# Seed functional ecology in Brazilian rock outcrop vegetation: an integrative synthesis

**DOI:** 10.1101/2023.03.21.533674

**Authors:** Carlos A. Ordóñez-Parra, Natália F. Medeiros, Roberta L.C. Dayrell, Soizig Le Stradic, Daniel Negreiros, Tatiana Cornelissen, Fernando A. O. Silveira

## Abstract

**Background and Aims:** Rock outcrop vegetation is distributed worldwide and hosts a diverse and unique flora that evolved under harsh environmental conditions. Unfortunately, seed ecology in such ecosystems has received little attention, especially regarding seed traits, germination responses to abiotic factors and the potential role of phylogenetic relatedness on such features Here, we provide the first quantitative and phylogenetically-informed synthesis of the seed functional ecology of Brazilian rock outcrop vegetation, with a particular focus on quartzitic and ironstone *campo rupestre*.

**Methods:** Using a database of functional trait data, we calculated the phylogenetic signal of seven seed traits for 371 taxa and tested whether they varied among growth forms, geographic distribution, and microhabitats. We also conducted meta-analyses that included 4,252 germination records for 102 taxa to assess the effects of light, temperature, and fire-related cues on the germination of *campo rupestre* species and explored how the aforementioned ecological groups and seed traits modulate germination responses.

**Key Results:** All traits and germination responses showed a moderate-to-strong phylogenetic signal. *Campo rupestre* species responded positively to light and had maximum germination between 20-25 °C. The effect of temperatures beyond this range was moderated by growth form, species geographic distribution, and microhabitat. Seeds exposed to heat shocks above 80 °C lost viability, but smoke accelerated germination. We found a moderating effect of seed mass for in responses to light and heat shocks, with larger, dormant seeds tolerating heat better but less sensitive to light. Species from xeric habitats evolved phenological strategies to synchronise germination during periods of increased soil water availability.

**Conclusions:** Phylogenetic relatedness plays a major role in shaping seed ecology of Brazilian rock outcrop vegetation. Nevertheless, seed traits and germination responses varied significantly between growth forms, species geographic distribution and microhabitats, providing support to the regeneration niche hypothesis and the role of functional traits in shaping germination in these ecosystems.

## INTRODUCTION

Rock outcrops are outstanding geological features where the bedrock protrudes above the land’s surface due to the erosion of softer parts of the landscape (Fitzsimons and Michael 2017). They offer a unique habitat that drastically contrasts with the neighbouring vegetation (Porembski 2007). Notably, they experience severe surface temperatures and have shallow, poorly-developed soils with low water retention capacity –a combination of abiotic factors that has driven the evolution of distinctive traits that allow plants to establish and survive in such harsh environments (Kluge and Brulfert 2000; Escudero *et al*. 2015; Oliveira *et al*. 2016; Bondi *et al*. 2023). Most studies, however, have focused on the ecophysiology of adult plants, and only few comprehensive reviews on germination ecology have been produced to date (e.g., Wyatt 1997; Biedinger et al. 2000). As a result, the potential role of regeneration traits (Donohue *et al*. 2010; Larson and Funk 2016) and regeneration niche (Grubb 1977) in shaping ecological processes in rock outcrops has been largely overlooked.

In Brazil, there are four main vegetation types associated with rock outcrops: *campo rupestre*, *canga*, *campo de altitude*, and inselbergs (Martinelli 2007). *Campo rupestre* occurs on quartzite, sandstone, and ironstone outcrops throughout the country, but it is most common at high elevations in the Espinhaço Range, eastern Brazil (Miola *et al*. 2021). This vegetation is characterised by sandy and rocky grasslands dominated by grasses, sedges and forbs (mostly Xyridaceae and Eriocaulaceae) with isolated patches of shrubs (mainly Melastomataceae, Fabaceae, Asteraceae, Myrtaceae, Vochysiaceae and Clusiaceae) and desiccation-tolerant and epilithic species establishing directly in the rock (Silveira *et al*. 2016). The *campo rupestre* that develops on ironstone outcrops receives the name *canga* and is functionally analogous to the quartzitic outcrops (Jacobi *et al*. 2007). The *campo de altitude* is found on granite and gneissic outcrops within the Atlantic Forest and consists of a mosaic of shrubs (mostly Asteraceae, Melastomataceae, and Myrtaceae) and small patches of dwarf forests amidst a matrix of grasses (Safford 1999). In contrast, inselbergs are isolated, dome-shaped granitic outcrops present across the country. In such geological formations, the mat-like vegetation –dominated mostly by Bromeliaceae, Cactaceae, Orchidaceae, Velloziaceae and shrubs from several families– is notably patchy, occurring only in depressions in the rock surface that accumulate soil (Safford and Martinelli 2000). All these open, grassy-shrubby, fire-prone and highly heterogeneous vegetation types are centres of species richness, phylogenetic diversity and endemism (Porembski 2007; Silveira *et al*. 2016; Campos *et al*. 2018).

The germination ecology of Brazilian rock outcrop vegetation has been previously reviewed (Garcia and Oliveira 2007; Nunes *et al*. 2016; Garcia *et al*. 2020), but these syntheses were restricted to the *campo rupestre* and focused on the differences in germination responses in a few emblematic families (e.g., Asteraceae, Bromeliaceae, Cactaceae, Eriocaulaceae, Melastomataceae, Velloziaceae, and Xyridaceae). Still, based on seminal studies on seed functional ecology, these syntheses put forward various promising hypotheses on the role of seed and germination traits in the ecological dynamics of plant communities. A first hypothesis poses that germination responses are shaped by seed traits, especially seed mass. For example, Nunes *et al*. (2016) pointed out that *campo rupestre* species with lighter seeds were more dependent on light for germination than larger ones, as expected under the trade-off between light requirements and seed mass (Milberg *et al*. 2000; Pearson *et al*. 2003). Still, such an association between seed traits and germination responses remains to be formally tested for this ecosystem. The second hypothesis is that contrasting microhabitat preferences and geographical ranges derive from distinct regeneration traits, as proposed by the regeneration niche hypothesis (Grubb 1977; Saatkamp *et al*. 2019). For instance, species from xeric microhabitats in the *campo rupestre* have been shown to germinate at higher temperatures than those from mesic environments (Oliveira and Garcia 2011), and broadly distributed species have broader germination niches than those restricted to a single substrate (Marques *et al*. 2014). Species with a broader distribution range have also been shown to have a wider germination temperature range (Ranieri *et al*. 2012). Similar studies, however, have failed to find such a significant association between microhabitat preferences or geographic distributions in the *campo rupestre* (Oliveira and Garcia 2011; Silveira et al. 2012; Giorni et al. 2018), suggesting the patterns might be more complicated than previously thought. Moreover, these studies have been limited to a handful of species within single families, making it impossible to draw broad conclusions for the entire rock outcrop community. A new opportunity to address these questions at the community level has emerged owing to a comprehensive database recently published (Ordóñez-Parra *et al*. 2023). Another limitation of previous reviews is that they did not explicitly address the potential role of shared evolutionary history in seed ecology. Local and global scale studies have highlighted that several seed traits, such as seed mass (Moles *et al*. 2005; Zanetti *et al*. 2020), dormancy (Willis *et al*. 2014; Dayrell *et al*. 2017) and germination requirements (Arène *et al*. 2017), are phylogenetically clustered, potentially due to a combination of adaptive evolution and phylogenetic conservatism (Vandelook *et al*. 2012; Carta, Mattana, *et al*. 2022). As a result, a proper vegetation-wide assessment of ecological hypotheses regarding seed ecology must incorporate the relative importance of phylogenetic relatedness in seed functional traits and germination responses.

Here, we conducted the first integrative synthesis of seed functional ecology of Brazilian rock outcrop vegetation to address the two previously described long-standing hypotheses and the potential role of phylogenetic relatedness on the seed ecology of these ecosystems. First, we assessed whether seed functional traits associated with different dimensions of the seed ecological spectrum exhibit a significant phylogenetic signal and evaluated whether they differ between growth forms, species geographic distribution and microhabitats. Second, we assessed the effect of light, temperature, and fire-related cues on the germination of plant species in these communities through phylogenetically informed meta-analyses. Such abiotic factors are established germination cues in these ecosystems and are the most studied in Brazilian rock outcrop vegetation, allowing us to provide more robust conclusions about their effects on germination (Figure S1). We then contrasted germination responses between growth forms, species geographic distribution and microhabitats to test the regeneration niche hypothesis (Grubb 1977). Finally, we tested if such responses were moderated by seed mass and dormancy –central traits associated with multiple seed functions (Saatkamp *et al*. 2019). Because rock outcrops occur in all continents and biomes, and their associated vegetation is quite similar in structure and function (Barthlott and Porembski 2000), we hope this study provides a solid starting point towards a global synthesis of the germination ecology of rock outcrop vegetation.

## MATERIALS AND METHODS

### Data sources

#### Seed functional traits and germination experiments

We retrieved seed trait and germination records from *Rock n’ Seeds* (Ordóñez-Parra *et al*. 2023), a database of seed functional traits and germination experiments of plant species from Brazilian rock outcrop vegetation (Figure 1). This database contains information on 16 functional traits in 383 taxa and 10,187 germination records for 281 taxa. We focused on seven functional traits associated with regeneration from seeds and major seed functions (Grubb 1977; Saatkamp *et al*. 2019): seed dry mass (mg), seed water content (%), percentage of embryoless seeds (%), percentage of viable seeds (%), seed dispersal syndrome (anemochory, autochory and zoochory), seed dispersal season (early-dry, late-dry, early-rain and late-rainy season) and primary dormancy [dormant (D) and non-dormant (ND)]. This subset of traits included data from 103 studies on 371 species. We also retrieved information from experiments assessing the effects of light availability, constant and alternate temperatures, and fire-related cues (i.e., heat shocks and smoke) from 22 studies and 102 species, totalising 4,252 germination records. Beyond seed and germination traits, the database also provides species-level data on growth form, microhabitat (‘xeric’ substrates experiencing pronounced seasonal drought, or otherwise ‘mesic’), and geographic distribution (‘restricted’ to rock outcrop vegetation, and ‘widespread’ if found in other vegetation types in addition to rock outcrops) (Ordóñez-Parra *et al*. 2023).

**Figure 1.**
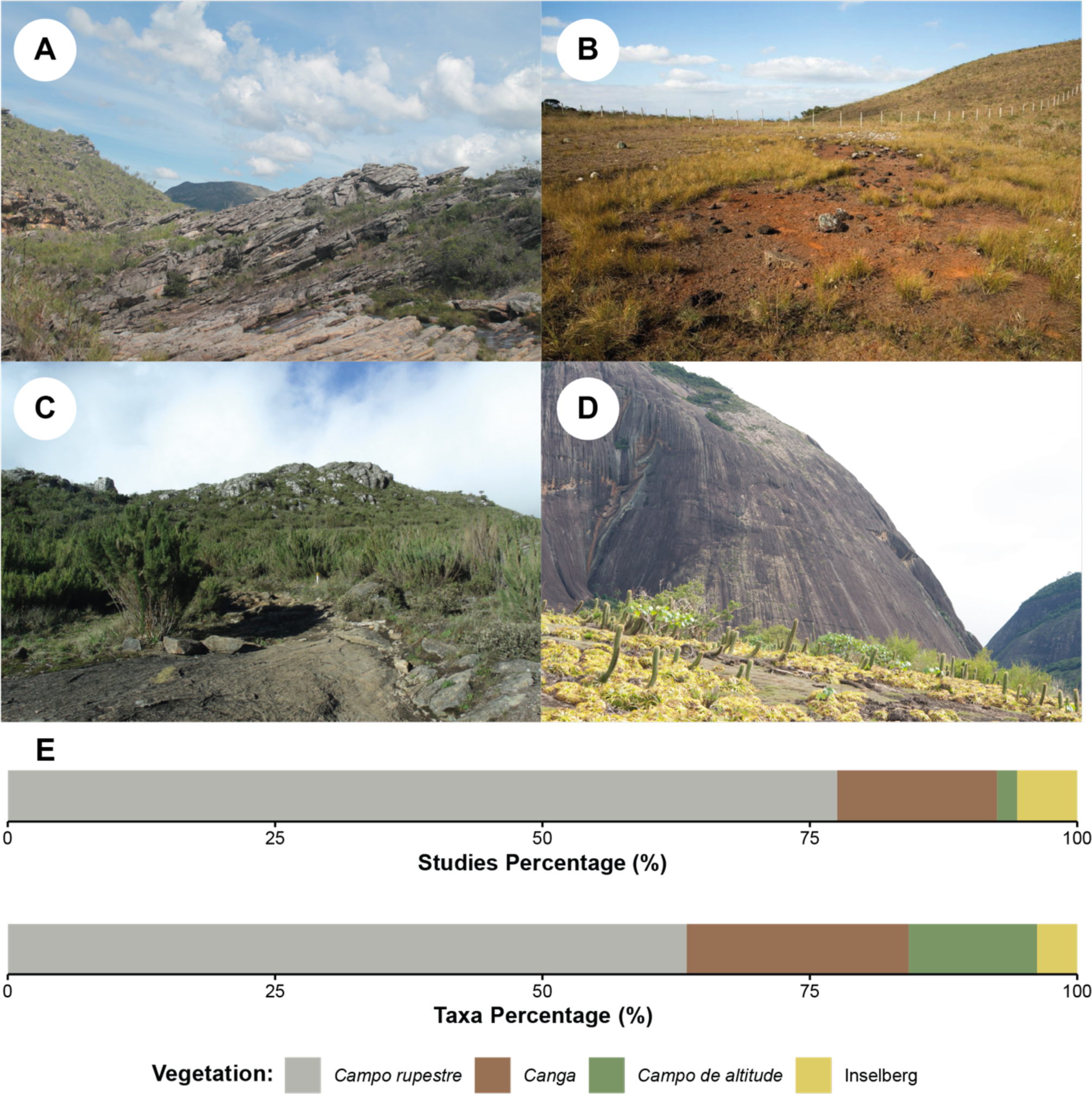
The four main vegetation types associated with rock outcrops in Brazil and its representation in the *Rock n’ Seeds* database (Ordóñez-Parra *et al*. 2023). A. A typical *Campo rupestre* landscape at Serra do Cipó. B. *Canga* vegetation at Parque Estadual da Serra do Rola Moça. C. *Campo de altitude* at Parque Nacional do Caparaó. D. Inselberg vegetation at Teófilo Otoni municipality. E. Percentage of studies and taxa evaluated on each vegetation type. All locations are in the State of Minas Gerais. Photos by Carlos A. Ordóñez-Parra (A), Roberta L. C. Dayrell (B), Daniela Calaça (C) and Fernando A. O. Silveira (D).

#### Phylogenetic tree

We reconstructed the phylogenetic tree of the studied species using the package *V.PhyloMaker2* (Jin and Qian 2022). First, species names were checked and updated following The Leipzig Catalogue of Vascular Plants using the R package *lcvplants* (Freiberg *et al*. 2020). We kept taxa identified at the species level, and all subspecies and varieties (18 cases) were upgraded to the species level. A phylogenetic tree for the resulting 370 species was generated, based on the GBOTB phylogeny (Smith and Brown 2018), which was updated and standardised following Freiberg *et al*. (2020). Taxa absent from this phylogeny were bound to their designated relatives using the bind.relative function of *V.PhyloMaker2* based on different sources (Almeda *et al*. 2016; Rivera *et al*. 2016). Species with no known relatives in the phylogeny (*Austrocritonia velutina*, *Cavalcantia glomerata, Cavalcantia percymosa* and *Parapiqueria cavalcantei*) were added using the at.node function from the *ape* package (Paradis and Schliep 2019). All these species belong to the Eupatorieae (Asteraceae), a tribe where infrageneric relationships remain unresolved (Rivera *et al*. 2016). Therefore, they were added to the base of the tribe as polytomies. Most infrageneric relationships in the phylogeny remained unresolved, appearing as polytomies, partially because the relationships within some genera of the highly diverse families in our database have low support (Alcantara *et al*. 2018; Guimarães *et al*. 2019). Nevertheless, phylogenetic metrics estimated from trees resolved to the genus level have shown to be highly correlated with those derived from fully resolved trees, suggesting that these polytomies do not weaken the results (Qian and Jin 2021).

### Statistical analyses

All analyses were made using R v. 4.3.2 (R Core Team 2023), and the code prepared will be provided as Supplementary Material and uploaded to GitHub upon acceptance (see Open Data).

#### Variation and phylogenetic signal of seed functional traits

To test for the phylogenetic signal in the quantitative seed traits (i.e., dry mass, water content, percentage of viable and embryoless seeds), we calculated Pagel’s λ (Pagel 1999) using the phylosig function from the package *phytools* (Revell 2012). λ values range from zero to one, with λ = 0 indicating that there is no phylogenetic signal; thus, related taxa are not more similar than expected by chance, and λ = 1 implying a strong phylogenetic signal, so the traits of related taxa are more similar than expected by chance (Pagel 1999). Pagel’s 11 was selected among the available phylogenetic signal indices given its robustness in measuring phylogenetic signals based on incompletely resolved phylogenies (Molina-Venegas and Rodríguez 2017). We also calculated Moran’s I coefficient to assess the phylogenetic autocorrelation of traits at the genus, family and order levels using the correlogram.formula function from *ape* (Paradis and Schliep 2019). Values of Moran’s I range from −1 to 1, with I = 0 indicating that species resemble each other as expected by the Brownian motion model. Values above and below zero imply that related species are more or less similar than expected by such model, respectively (Moran 1950; Gittleman and Kot 1990). All these tests were carried out using log-transformed seed mass values and logit-transformed water content, percentage of embryoless and viable seed values to reduce the skewness of the data. For species with more than one record (25 species), we carried out the analyses using the average species value.

For the qualitative seed traits (i.e., seed dormancy, seed dispersal syndrome and seed dispersal season), the phylogenetic signal was tested by implementing the rtestdeciv function from the *adiv* package using 9,999 permutations (Pavoine 2020). This method decomposes trait diversity among the nodes of the phylogenetic tree and assesses whether it is skewed towards the tree’s root or tips, with significant root skewness implying the presence of a significant phylogenetic signal. Seed dormancy and dispersal season were treated as multichoice variables, as 11 species had records of both D and ND seeds, and 18 species reported more than one dispersal season.

To test for differences in quantitative traits between ecological groups (growth form, microhabitat, and geographic distribution), we used phylogenetic generalised least square models as implemented in the package *caper* (Orme *et al*. 2018). Likewise, qualitative trait variation was assessed using a phylogenetic logistic regression (Ives and Garland 2010) implemented in the *phylolm* package (Ho and Ané 2014). Because dispersal season and syndrome had more than two possible states, individual models were run to assess the probability of each season and syndrome. Species with records for more than one distribution class were classified as “widespread”, and species with occurrence in both microhabitats were coded as “mesic/xeric”. For this analysis, species with records from more than one combination of qualitative traits were included as different populations, using the add.taxa.phylo function from the *daee* package (Debastiani 2021) and keeping the tree ultrametric.

Preliminary tests showed that the interaction between our qualitative predictors did not significantly affect any of the tested variables. Moreover, the models that included such interactions were outperformed by simple additive models based on Akaike Information Criteria (see code provided). As a result, our final models did not include these interactions. Growth form comparisons were made between herbs and shrubs only due to the low sample size of trees (26 species), succulents (nine species) and lianas (six species).

#### Meta-analyses of germination responses to abiotic factors

We estimated the overall effects of i) light, ii) constant and alternate temperatures, and iii) fire-related cues on germination through meta-analyses. We achieved this by fitting phylogenetic generalised linear mix models with Bayesian estimations using the Markov chain Monte Carlo (MCMC) method –an approach that has been previously used in meta-analyses of primary germination data (Vandelook et al. 2018; Fernández-Pascual et al. 2021; Fernández-Pascual et al. 2022; Carta, Fernández-Pascual, et al. 2022). This analysis was restricted to *campo rupestre* and *canga* species, given the lack of data from *campo de altitude* and the small sample size of independent observations for inselbergs (two studies, three species) for these abiotic factors. Germination records under complete darkness were used as controls to test the effects of light. Also, given the overall positive effect of light on germination (see Results), only experiments carried out under light conditions were used to assess the effects of different temperatures and fire-related cues on seed germination. We used 25 °C as a control for constant and alternate temperatures because it is considered the optimum temperature for most species in this ecosystem (Garcia and Oliveira 2007; Nunes *et al*. 2016; Garcia *et al*. 2020). Finally, we used untreated seeds (i.e., seeds not exposed to either heat shock or smoke) as controls for fire-related cues.

We assessed the effects of these abiotic factors on the proportion of germinated seeds and median germination time (t_50_). The number of germinated and ungerminated seeds was taken from the raw germination data deposited in *Rock n’ Seeds*. To avoid confounding effects from treatments with extremely low germination, we only included observations where either the control or treatment had ≥10% germination. The t_50_ (Coolbear *et al*. 1984 as modified by Farooq *et al*. 2005) was calculated for the experiments that included raw daily germination data using the *germinationmetrics* package (Aravind *et al*. 2022). We calculated this index only for observations where both the control and treatment conditions had ≥10% germination. Moreover, we assessed the effect of light, constant and alternate temperatures as germination cues in ND seeds or in seeds where dormancy has been experimentally alleviated. Alternate temperatures are known to break physical dormancy (Baskin and Baskin 2014), but none of the studies in the dataset have tested this. In contrast, we included D species in the meta-analyses of fire-related cues because these are known to break the seed dormancy in some species in fire-prone ecosystems (Baskin and Baskin 2014). Species with non-conclusive dormancy records were excluded.

We evaluated the effect of each abiotic factor (i.e. light, temperature and fire) on germination proportions and t_50_ on the whole dataset and separately for each growth form (herbs and shrubs), geographic distribution (widespread and restricted) and microhabitat (mesic, xeric, and mesic/xeric). We also tested the interaction of seed mass and seed dormancy on germination responses in the whole dataset, and the interaction effect of seed dormancy only for responses to fire-related cues. Phylogeny, species identity, study and observations were included as random variables. Because seed mass exhibited a significant, strong phylogenetic signal (see Results), the missing seed mass values for 54 species were inputted using average values for the genera from *Rock n’ Seeds* (Ordóñez-Parra *et al*. 2023) and the Seed Information Database (https://ser-sid.org/). The analyses were performed with the MCMCglmm package (Hadfield 2010). All models were run with weakly informative priors, with parameter-expanded priors for the random effects. Each model was run for 500,000 MCMC steps, with an initial burn-in phase of 50,000 and a thinning interval of 50 (de Villemereuil and Nakagawa 2014), resulting in an average of 9000 posterior distributions. We calculated mean parameter estimates and 95% credible intervals (CIs) from the resulting posterior distributions. CIs overlapping zero were considered non-significant. To estimate the phylogenetic signal of germination responses, we also estimated the Pagel’s λ of each model, as described by de Villemereuil and Nakagawa (2014).

## RESULTS

### Phylogenetic signal and variation of seed functional traits

All examined seed traits exhibited a significant phylogenetic signal (Figure 2, Table 1, *p* < 0.001). For all quantitative traits (i.e., seed mass, water content, percentage of embryoless and percentage of viable seeds), the phylogenetic signal was moderate to strong (11 between 0.57-0.90), meaning that closely related taxa strongly resembling each other both at the genus and family level (Table S1). At the order level, we only found a significant and negative phylogenetic autocorrelation for seed mass (I = −0.39, *p* < 0.001, Table S1). Our data did not suggest the presence of phylogenetic autocorrelation for seed water content and the percentage of embryoless and viable seeds at the order level (*p* > 0.05). A summary of family-level variation in quantitative and qualitative traits is presented in Figure S2.

**Figure 2.**
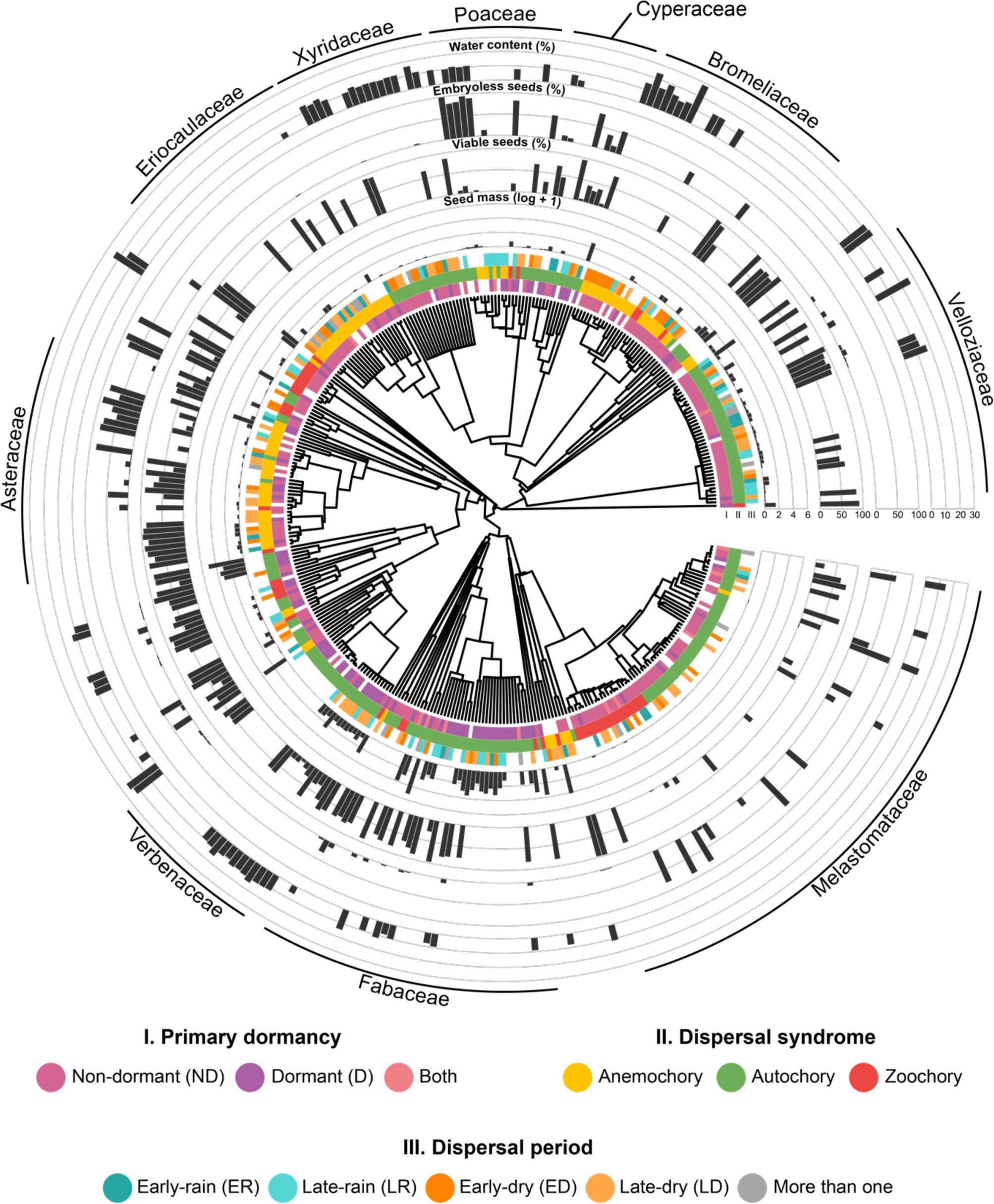
Phylogeny of studied species with available information on seed functional traits. Bars represent trait values for each species. The ten families with the largest number of species in the dataset are labelled. Figure elaborated with the R packages ggtree (Yu *et al*. 2017), ggtreeExtra (Xu *et al*. 2021) and ggnewscale (Campitelli 2022).

**Table 1.** Results of phylogenetic signal test for seed traits in Brazilian rocky outcrop vegetation. P-values for quantitative traits come from the likelihood ratio test performed by phylosig function, while it corresponds to the root-to-tip skewness test performed by the rtestdecdiv function for qualitative traits.

| Quantitative traits | Lambda | p-value |
| --- | --- | --- |
| Seed mass | 0.90 | < 0.001 |
| Water content | 0.72 | < 0.001 |
| Percentage of embryoless seeds | 0.75 | < 0.001 |
| Percentage of viable seeds | 0.57 | < 0.001 |
| Qualitative traits | p-value |  |
| Primary dormancy | < 0.001 |  |
| Dispersal syndrome | < 0.001 |  |
| Dispersal season | < 0.001 |  |

Regarding the variation of quantitative traits, seed mass variation differed between growth forms and microhabitats, with shrubs (t = 2.19, p = 0.030) and species inhabiting both mesic and xeric microhabitats (t = 2.01, p = 0.045) producing heavier seeds than herbs and species restricted to xeric microhabitats, respectively. Seed water content, percentage of embryoless and viable seeds did not vary between any of the groups, and seed mass did not differ between species with distinct geographic distributions (Table 2).

**Table 2.** Differences in seed functional traits between growth forms (herbs vs. shrubs), and species distribution (restricted vs. widespread) and microhabitat (mesic vs. mesic/xeric vs. xeric) in Brazilian rocky outcrop vegetation. Bold values indicate p-values < 0.05. t and z correspond to the values obtained in the t- and z-tests that are part of the implementation of phylogenetic least square models and phylogenetic logistic regressions, respectively.

|  | Growth form: |  | Distribution: |  | Microhabitat: |  | Microhabitat: |  |
| --- | --- | --- | --- | --- | --- | --- | --- | --- |
|  | Shrub |  | Widespread |  | Mesic/Xeric |  | Xeric |  |
| Quantitative traits | t | p-value | t | p-value | t | p-value | t | p-value |
| Seed mass | **2.1872** | **0.0298** | -0.5986 | 0.5501 | **2.0141** | **0.0453** | 1.6115 | 0.1086 |
| Water content | 0.2789 | 0.7811 | -0.0770 | 0.9388 | 0.6075 | 0.5453 | 0.7834 | 0.4358 |
| Embryoless seeds | 1.9274 | 0.0572 | -0.2368 | 0.8134 | 0.9752 | 0.3322 | 1.2967 | 0.1982 |
| Viable seeds | -1.0221 | 0.3085 | 0.2381 | 0.8122 | -0.3040 | 0.7616 | 0.7006 | 0.4847 |
| Qualitative traits | Z | p-value | z | p-value | z | p-value | z | p-value |
| Dormancy | 1.4125 | 0.1578 | 1.5052 | 0.1323 | 1.5266 | 0.1268 | **2.5218** | **0.0117** |
| Anemochory | -1.2154 | 0.2242 | 1.3943 | 0.1632 | 1.0047 | 0.3150 | 1.1497 | 0.2503 |
| Autochory | 0.0387 | 0.9691 | -0.0710 | 0.9434 | -0.0792 | 0.9368 | -0.0998 | 0.9205 |
| Zoochory | 0.3290 | 0.7422 | 0.0048 | 0.9962 | 0.6844 | 0.4937 | -0.0040 | 0.9968 |
| Early-Dry | -0.9086 | 0.3636 | -0.1266 | 0.8993 | -1.0427 | 0.2971 | 0.4504 | 0.6524 |
| Late-Dry | **3.4323** | **0.0006** | -0.7443 | 0.4567 | 0.7421 | 0.4580 | -1.3673 | 0.1713 |
| Early-Rain | 0.1036 | 0.9175 | -0.7541 | 0.4508 | -0.4839 | 0.6284 | 0.0445 | 0.9645 |
| Late-Rain | **-2.0137** | **0.0440** | -0.1826 | 0.8551 | **2.0319** | **0.0422** | **2.1507** | **0.0315** |

For qualitative traits, we found that shrubs had a higher probability of dispersing seeds during the late-dry season (z = 3.43, p < 0.001), while dispersal during the late-rainy season was more likely in herbs (z = −2.01, p = 0.044). Primary dormancy and dispersal season varied significantly between microhabitats, with species from xeric environments having higher probabilities of producing dormant seeds (z = 2.52, p = 0.012) and dispersing their seeds during the late-rainy season than species from mesic microhabitats (z = 2.15, p = 0.031). Similarly, species from both mesic and xeric microhabitats also had a higher probability of dispersing seeds during the late-rainy season (z = 2.03, p = 0.042). We found no significant differences in the probability of any of the dispersal syndromes or the other dispersal seasons (Table 2).

In summary, shrubs tended to produce heavier seeds and to have higher probabilities of dispersal during the late-dry season. In contrast, herbs produced relatively smaller seeds with higher probabilities of dispersal during the late-rainy season. Species from mesic/xeric microhabitats tended to produce heavier seeds and disperse them during the late-rainy season compared to those exclusively from mesic microhabitats. Moreover, species restricted to xeric environments had higher probabilities of producing dormant seeds during late-rainy dispersal than those from mesic environments. Finally, species restricted to outcrop vegetation and widespread ones did not differ in any seed trait (Figure 3).

**Figure 3.**
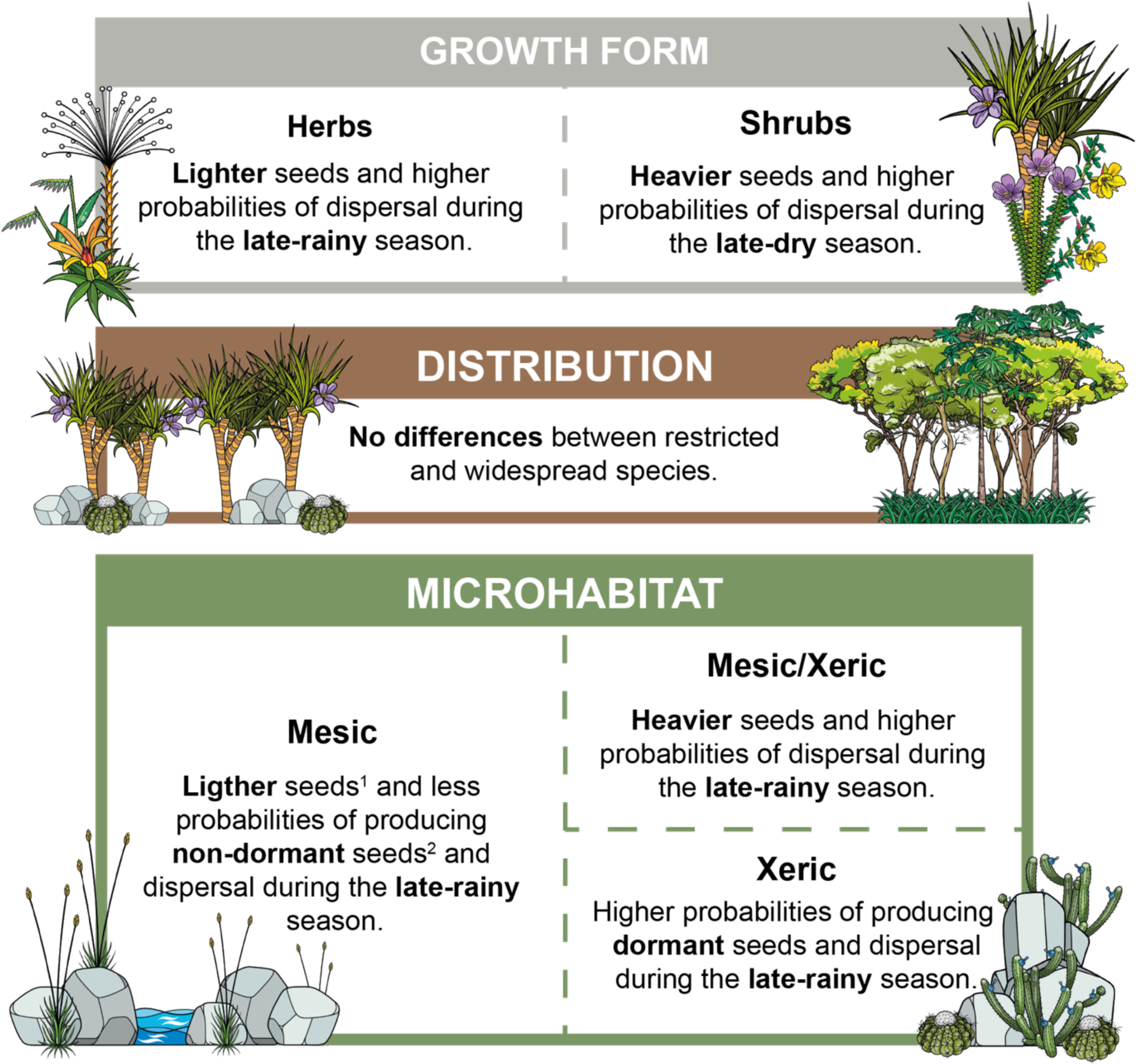
Summary of seed trait differences among growth forms, species geographic distribution and microhabitat, based on results from Table 2. Notes: 1. When compared to species form mesic/xeric microhabitats. 2. When compared to species from xeric microhabitats.

### Meta-analyses of germination responses

Light positively and strongly affected the final germination proportion regardless of growth form, distribution, or microhabitat. Still, light effects were higher for shrubs and species from mesic environments compared to herbs and species from other microhabitats, respectively. Seed responses to light did not vary between species with different geographic distributions. Seed mass had a negative interactive effect with responses to light (Figure 4).

**Figure 4.**
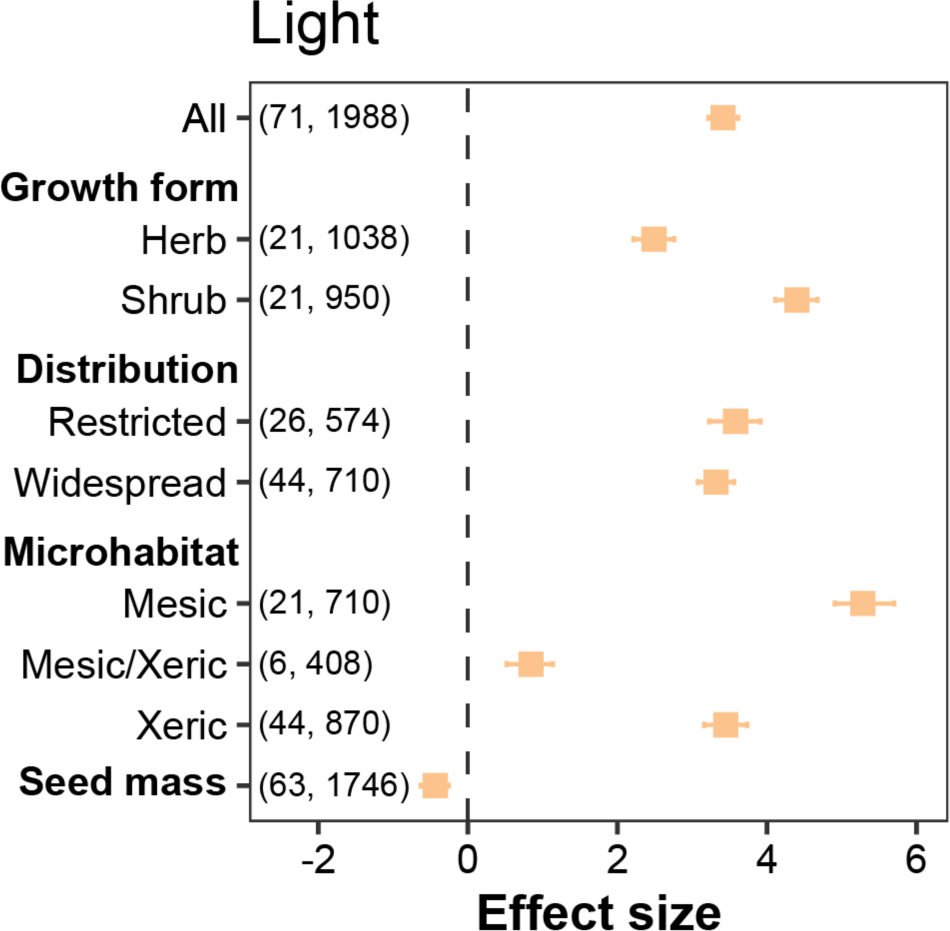
Effect of light availability on germination proportion in *campo rupestre* species (Control: total darkness). Squares indicate the posterior means of the effect for each moderator (growth form, distribution and microhabitat) and whiskers the 95 % credible intervals of the effect size. Numbers in parentheses indicate the number of species and observations (each observation is a replicate of a given germination experiment) employed for each model.

Regarding the overall effect of constant temperatures on germination, temperatures below 20 °C had no significant effect on germination proportions when compared to the control. However, temperatures below and above 20 °C had a consistent negative effect on germination proportion and a positive effect on t_50_. Herbs and shrubs exhibited differential responses to temperature regimes: germination proportion decreased more with decreasing temperature for shrubs, whereas the final germination proportion of herbs increased at 20 °C and remained relatively constant at 30 °C. For instance, the t_50_ of herbs was not significantly affected by 30 °C, contrasting with shrubs for which germination was slowed at this temperature. Differential responses were also found for species with contrasting geographic distributions, with species with widespread distribution showing a more severe reduction in the germination proportion at 10 °C as well as a significant negative reduction at 30 °C. Similarly, the germination proportion of species from mesic microhabitats was less negatively affected by 15 °C conditions but was more inhibited at 35 and 40 °C than species from xeric microhabitats. Finally, seed mass exhibited a significant interaction with germination responses to temperature, being positive for germination proportion at 10 and 35 °C and negative at 20 and 40 °C. Still, we did not find a significant interaction between seed mass and germination responses at 15 and 30 °C (Figure 5A).

**Figure 5.**
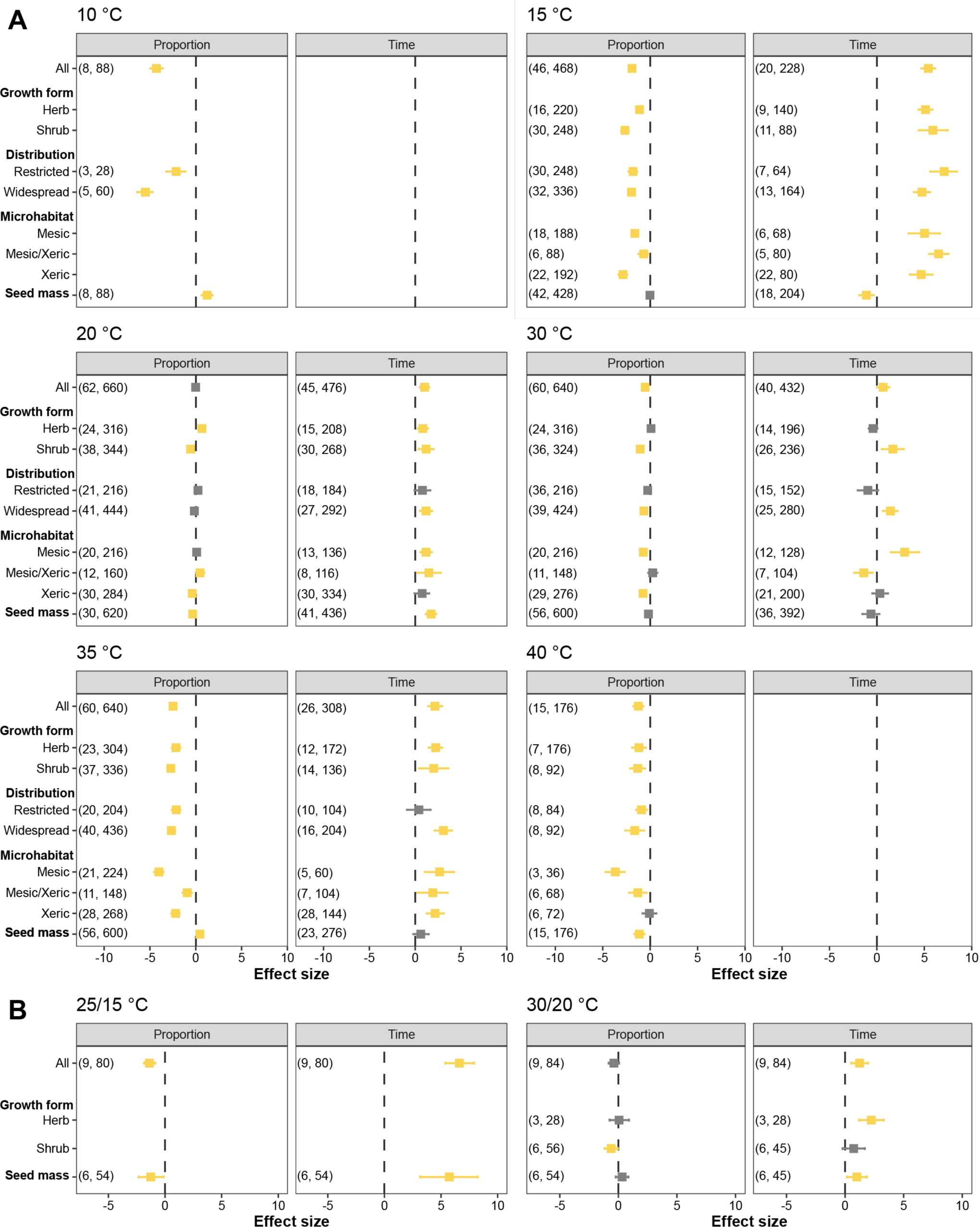
Germination responses to (A) constant and (B) alternate temperatures in *campo rupestre* species (Control: 25 °C). Squares indicate the standardised mean effect size for each moderator (growth form, distribution and microhabitat) and whiskers the 95 % confidence interval of the effect size. Coloured estimates indicate significant effects (i.e., where confidence intervals do not overlap zero). Numbers in parentheses indicate the number of species and observations (each observation is a replicate of a given germination experiment) employed for each model. The empty panel for germination time at 10 and 40 °C refers to a lack of data.

When evaluating the effects of alternate temperatures, we found that 25/15 and 30/20 °C regimes had an overall negative and a non-significant effect on germination proportion, respectively. Moreover, both treatments significantly increased t_50_. 30/20 °C regime also had a negative effect on the germination proportion in shrubs and increased t_50_ in herbs. Seed mass had a negative interaction with germination proportion at 25/15 °C and a positive interaction with t_50_ at both treatments (Figure 5B).

Finally, we found that fire-related cues reduced germination proportion while also affecting t_50_. The effect varied according to the treatment applied, with heat shocks –alone or in combination with smoke– reducing germination proportion but not germination time. Instead, smoke alone or in combination with heat did not significantly affect germination proportion but significantly reduced t_50_ (Figure 6A). When comparing heat shock treatments, we found that treatments heat shocks of 100 (both of 1 and 5 minutes) and 200 °C for 1 minute significantly reduced germination proportion, although they did not affect t_50_ (Figure 6B). This negative effect of heat shocks on germination proportion varied between growth forms, species distribution and microhabitats, with only shrubs, restricted species and species from xeric microhabitats being negatively affected by heat shocks. Instead, heat shocks did not significantly affect herbs, widespread species, and species from mesic microhabitats (Figure 6C). The negative effect of smoke on t_50_ remained significant for shrubs but was non-significant in herbs. Our analysis did not indicate that smoke’s effect was modulated by seed mass (Figure 6D). Contrastingly, primary dormancy and seed mass altered germination responses to heat shocks, with ND seeds exhibiting a reduction in germination proportion and seed mass holding a positive effect (Figure 6C).

**Figure 6.**
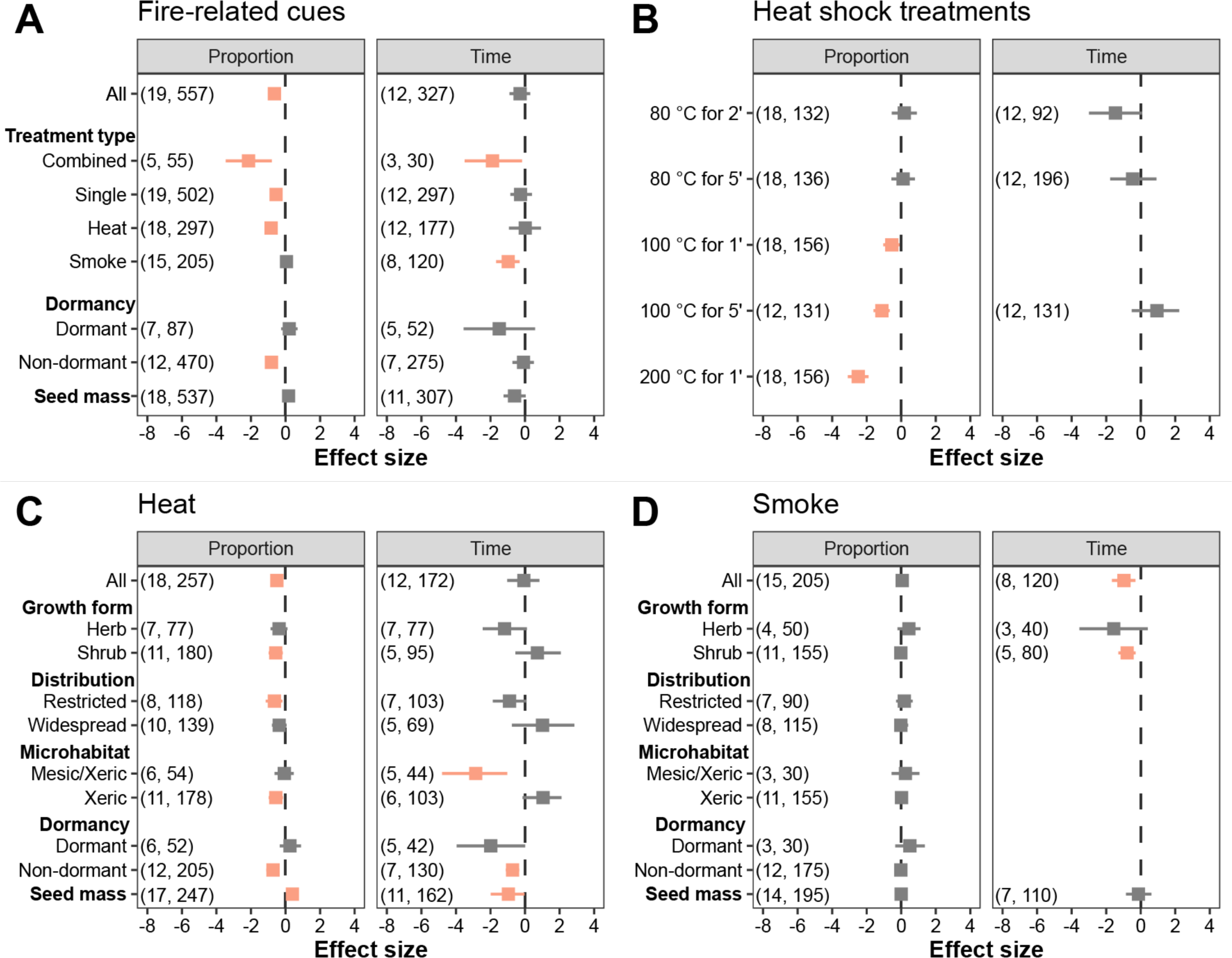
Germination responses to fire-related cues in *campo rupestre* species (Control: untreated seeds). (A) Overall effects. (B) Heat shocks temperatures. (C) Heat shocks without the 200 °C for one minute. (D) Smoke. Squares indicate the standardised mean effect size for each moderator (growth form, distribution and microhabitat) and whiskers the 95 % confidence interval of the effect size. Coloured estimates indicate significant effects (i.e., where confidence intervals do not overlap zero). Numbers in parentheses indicate the number of species and observations (each observation is a replicate of a given germination experiment) employed for each model.

Overall, phylogeny played a major role in germination responses to all these abiotic factors, with most of our models having a λ significantly different form zero. Moreover, in most cases, species identity had a stronger effect on germination responses variability than study or observations within studies (Table S2 and S3).

## DISCUSSION

### Seed mass shapes germination responses in campo rupestre

Our analysis points out a significant interaction between seed mass and germination responses to light, constant temperature and heat shocks, supporting the long-standing first hypothesis that seed mass modulates germination responses. Regarding light, we found a significant and negative interaction between seed mass and germination responses to light, with larger seeds being less responsive to light availability. Such results support the hypothesis raised by Nunes *et al*. (2016) on the existence of this trade-off in the *campo rupestre* (Milberg *et al*. 2000; Pearson *et al*. 2003).

We also found that seed mass had a significant interaction with response to some temperatures. Nevertheless, there was no overall pattern between germination responses to temperature and seed mass, with a positive interaction with changes in germination proportion at 10 and 30 °C, a negative one at 20 and 40 °C and non-significant interactions at 15 and 35 °C. Despite being significant, the effect of such interactions is rather small (see Table S2 and S3), which agrees with the results of Arène *et al*. (2017), who found a weak association between cardinal temperatures and seed mass that disappeared when accounting for phylogenetic relatedness. Therefore, it seems reasonable to conclude that variation in germination responses in *campo rupestre* is probably better explained by phylogeny rather than by seed mass, an observation supported by the high λ values found in our models. Moreover, the small seed mass variation in our dataset (86% of records < 1 mg) could prevent us from finding a more consistent pattern for the role of seed mass in this ecosystem.

We also found a significant interaction between seed mass and germination responses to heat shocks that indicate that *campo rupestre* species with larger seeds tend to have increased and faster germination after a heat shock, a result that is consistent with a positive correlation between seed mass and heat tolerance reported in the Cerrado (Ribeiro *et al*. 2015; Daibes *et al*. 2019). Such association can be explained by decreasing surface-to-volume ratio with increasing seed mass, indicating that larger, heavier seeds have higher insulation against higher, lethal temperatures (Ruprecht *et al*. 2015).

Finally, the lack of a significant effect of alternate temperatures or an interaction of such responses with seed mass could be explained by the fact that most *campo rupestre* species produce small, lighter seeds, which are not expected to benefit from alternate temperature regimes (Pearson *et al*. 2003).

### Seed functional traits are phylogenetically clustered in Brazilian rock outcrop vegetation

Our results indicate that all seed traits showed moderate-to-strong phylogenetic signals, which is consistent with global studies showing phylogenetic clustering for multiple seed traits, such as seed mass (Moles *et al*. 2005) and dormancy (Willis *et al*. 2014). However, our results partially contrast with local studies in similar ecosystems. First, Zanetti *et al*. (2020) did not find a phylogenetic signal for seed water content in their study of 48 species from the *cangas* of Carajás (Eastern Brazilian Amazon). This discrepancy between this and our study might arise from differences in sample size and geographic extent or the distinct phylogenetic structure of each rock outcrop vegetation type (Massante *et al*. 2023). Second, we found a significant phylogenetic signal for seed dispersal season, while seed dispersal season in the Cerrado (Escobar *et al*. 2021) and other phenological events in the *campo rupestre* (Zanetti *et al*. 2020; Oliveira *et al*. 2021) have not shown such signal. As a result, seed dispersal season is the only phenophase that has shown a significant phylogenetic signal in the Brazilian rock outcrop vegetation, potentially because of an intense evolutionary pressure towards germination timing in these harsh ecosystems, where water is even scarcer than in the Cerrado (Oliveira et al. 2021), thus may create stronger selective pressures for establishment.

Seed mass and seed dispersal season differed between herbs and shrubs, with shrubs tending to produce relatively larger seeds during the late-dry season and herbs producing lighter seeds during the late-rain season. Differences in seed mass between herbs and shrubs have been attributed to the distinct strategies employed by such growth forms: while shrubs invest in fewer, larger seeds that cope better with environmental hazards, herbs invest in numerous, lighter seeds to increase their establishment opportunities (Westoby *et al*. 2002; Moles *et al*. 2005). This relationship is also supported by the global spectrum of plant form and function (Díaz *et al*. 2016), suggesting that this global pattern also holds at small scales, such as for Brazilian rock outcrop vegetation. On the other hand, differences in seed dispersal season suggest differences in phenological strategies to cope with precipitation seasonality (see below).

### Species from distinct geographic distributions and microhabitats exhibit different germination responses (and seed traits)

Rock outcrops and the surrounding vegetation differ in several aspects that should impact germination, such as nutrient and water availability, irradiance, and daily temperature fluctuations (Oliveira *et al*. 2016). While we did not find significant differences in seed traits between restricted and widespread species, their germination responses to temperature and heat shocks differed. First, the germination proportion of widespread species was reduced or delayed at relatively low and high temperatures, implying they are more susceptible to these conditions than species restricted to outcrop vegetation. Second, species restricted to outcrop vegetation were more sensitive to heat shocks than widespread ones. The rocky substrate of the *campo rupestre* does not favour fire spread (Conceição *et al*. 2016), so natural fire events in these ecosystems are unlikely to reach the vegetation establishing directly on rock outcrops, reducing evolutionary pressures towards heat tolerance. In contrast, widespread species present in our dataset include species from the Cerrado, where fire events are relatively more frequent, and seeds exhibit relatively higher heat tolerance (Daibes *et al*. 2022).

Contrastingly, species from mesic and xeric microhabitats were found to differ both in terms of seed traits and germination responses, agreeing with local studies showing a correspondence between seed traits and species microhabitats (Oliveira and Garcia 2011; Marques et al. 2014; Ranieri et al. 2012). These results support our second hypothesis that predicts distinct germination requirements expected between species with contrasting microhabitat preferences and geographical ranges, as expected under the regeneration niche hypothesis (Grubb 1977). Interestingly, these results contrast with large-scale studies carried out in temperate ecosystems where an association between ecological preferences and germination responses has been found (Fernández-Pascual *et al*. 2021; Fernández-Pascual *et al*. 2022), suggesting that the extreme abiotic conditions of both the *campo rupestre* and rock outcrop vegetation as a whole impose a stronger selective pressure to germination traits. However, seed traits did not differ in species with distinct geographic ranges suggesting that multiple pressures operate on the selection of seed traits at larger scales.

Regardless of microhabitat, most species in our dataset dispersed their seeds during the late-or the early-dry season. However, species from xeric habitats had higher probabilities of producing dormant seeds and dispersing their seeds during the late-rain season –two strategies presumably arising due to evolutionary pressures towards strategies to synchronise germination with optimum conditions for seedling establishment, that is under higher soil water availability (Jurado and Flores 2005; Silveira, Ribeiro, *et al*. 2012; Garcia *et al*. 2020). Nevertheless, only 36% of our species produce dormant seeds, and *campo rupestre* is known to have the highest ND:D ratio globally (Dayrell *et al*. 2017), implying that seed dormancy might not be the main driver of seedling establishment in our study system. Additional strategies to control germination timing include germination requirements and the acquisition of secondary dormancy. A mismatch between temperature germination requirements and environmental conditions at the moment of dispersal might provide an alternative strategy to prevent germination in the absence of dormancy (Escobar *et al*. 2021) –for example, seeds dispersed during the dry season have evolved to germinate under relatively higher temperatures of the rainy season. Furthermore, secondary dormancy, as reported in a few Eriocaulaceae and Xyridaceae species from *campo rupestre* (see Garcia *et al*. 2020), can contribute to fine-tuning seedling establishment with optimum conditions. All these species disperse their non-dormant seeds between the early-dry and the early-rain season and become increasingly dormant as the rainy season advances. When the dry season starts, dormancy is progressively alleviated by reduced temperatures and low water availability, allowing germination at the onset of the rainy season (Duarte and Garcia 2015).

Species from mesic microhabitats also exhibit distinct responses to light and temperature, with stronger positive responses to light and being more tolerant to low (below 20 °C) but more sensitive to high temperatures (above 30 °C). As a result, a stronger response to light could ensure that germination under light conditions occurs as quickly as possible. Moreover, high soil humidity in mesic habitats buffers these habitats from wide temperature fluctuations and contributes to maintaining relatively low soil temperatures (Oliveira and Garcia, 2011).

### Campo rupestre species depend on light for germination

Our meta-analysis showed that light positively affects seed germination across all ecological groups, supporting previous assessments about its importance in *campo rupestre* germination ecology. Interestingly, we also found that germination responses to light were stronger in shrubs, a result that could be attributed to the fact that species with the lightest seeds in our dataset are shrubs from the Melastomataceae (see Figure S2), which is both the best-represented family in our dataset (Ordóñez-Parra *et al*. 2023) and a family characterised by an absolute light requirement for germination (Ordóñez-Parra *et al*. 2022).

Small-seeded species from our study are expected to have narrowly defined microsite requirements due to their limited internal resources, so further aspects of the light environment, such as spectral quality and the interaction between light and temperature, are expected to control their germination (Pearson *et al*. 2003; Pons 2014). Still, studies assessing such aspects in our study system are scarce and have only involved very few species (Pereira *et al*. 2009; Hmeljevski *et al*. 2014; Vieira *et al*. 2018; Garcia *et al*. 2020), preventing robust inferences about the functional relevance of these germination cues.

### Herbs and shrubs respond differently to low and high temperatures

We found that the germination of *campo rupestre* species was maximised between 20-25 °C, supporting previous qualitative reviews (Nunes *et al*. 2016; Garcia *et al*. 2020). Decreased germination proportion below this range could be a putative mechanism to avoid germination during the dry season when temperatures decrease and soil water potential does not support seedling establishment (Garcia *et al*. 2020). The negative effect of low temperatures was stronger in shrubs, suggesting that these require higher temperatures to germinate. In contrast, herbs had increased germination at 20 °C but were unaffected by 30 °C conditions. These results suggest that herbs dominate microsites experiencing lower soil temperatures.

Alternate temperature regimes had either a negative or null effect on the germination. The reduction of germination proportion and the increase of germination time at 25/15 °C regime is probably due to part of these temperatures being suboptimal. Conversely, studies in the *campo de altitude* have shown that several species benefit from alternate temperature regimes, suggesting this cue could be more important in this particular vegetation that occurs on higher elevation belts (Andrade *et al*. 2021). Therefore, additional studies are required to elucidate the functional relevance of alternate temperatures and whether this varies between vegetation types.

### Heat kills small, non-dormant seeds, but smoke accelerates germination

Seeds exposed to 100 or 200 °C heat shocks had their germination proportion reduced, while milder heat shocks (≤ 80 °C) did not affect germination proportion or time, supporting studies showing heat tolerance in the Cerrado (Daibes *et al*. 2022). In addition to seed mass, responses to heat shocks were also moderated by seed dormancy, with ND seeds being relatively less tolerant to heat shocks, as previously described for Cerrado species (Ramos *et al*. 2016; Daibes *et al*. 2019). Dormant species were mostly unaffected by heat shocks, supporting the notion that fire-mediated dormancy alleviation is not expected in rocky areas or savannas (Pausas and Lamont 2022). The lack of detrimental effects on dormant species also implies that traits associated with seed dormancy promote heat tolerance, such as the accumulation of heat shock proteins or the presence of hard and water-impermeable coats (Tweddle *et al*. 2003; Ramos *et al*. 2016).

Smoke did not affect germination proportion but accelerated germination, supporting similar results found for Cerrado vegetation (Motta *et al*. 2024) and the notion that smoke-derived compounds stimulate germination of ND seeds or those where dormancy has been alleviated (Mackenzie *et al*. 2021). Smoke-stimulated germination in *campo rupestre* is thought to promote the germination of species resprouting and shedding seeds after fire (Fernandes *et al*. 2021) –a usual phenological syndrome in this vegetation (Figueira *et al*. 2016)– allowing recently-dispersed seeds to exploit the post-fire environment where competition is relaxed (Le Stradic *et al*. 2015; Fernandes *et al*. 2021). While our meta-analysis supports this hypothesis, we found that the hastening effect of smoke is minor and only significant for shrubs. Moreover, such an effect has only been tested on a handful of species, and further studies are needed to rigorously test it.

## Conclusion

Seed germination ecology in rock outcrop vegetation in Brazil is strongly shaped by species phylogenetic relatedness, as illustrated by the moderate-to-strong phylogenetic signal exhibited across seed traits and germination responses. As a result, knowledge about one species can provide highly relevant information about the germination ecology of other closely related species. Despite that, our analyses also point out that seed mass can shape germination responses that may differ between species with different growth form, species geographic distribution and microhabitats. Our integrative synthesis provides robust support for long-standing hypotheses in the seed ecology of Brazilian rock outcrop vegetation. We hope that our synthesis paves the way for future research about the functional role of seed traits in these megadiverse ecosystems and a solid baseline for using seeds in ecological restoration and biological conservation programs, such as direct seeding and seed banking.

## ACKNOWLEDGEMENTS

The first author dedicates this article to the memory of Martha Galeano (1964-2022), who was fascinated with *tepuis* and whose Vegetation Studies lecture inspired him to pursue a career in plant ecology. This study is part of the first author’s Master Dissertation at the Plant Biology Program at Universidade Federal de Minas Gerais. CAO-P and NFM were supported by a scholarship from CAPES and CNPq, respectively. FAOS acknowledges support from FAPEMIG. We also thank James Dalling and Sergey Rosbakh, who provided valuable comments to an earlier version of the manuscript, and Rohit Goswami, who helped in elaboration of the figures and the organization of the GitHub repository.

## OPEN DATA

The R code for the analysis and creation of the figures will be provided as Supplementary Material and uploaded to GitHub upon acceptance.

---

**Figure S1.**
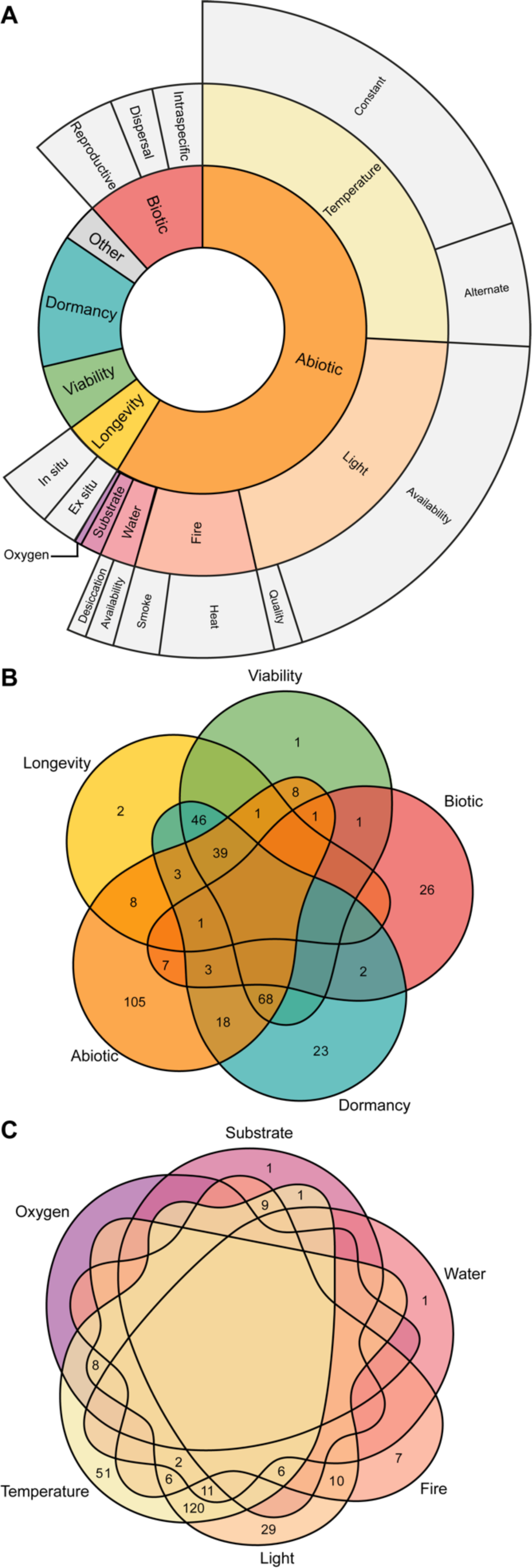
State of the art of the germination ecology of Brazilian rock outcrop vegetation. A. Percentage of studies for each germination ecology topic. B. Number of species studied each major topic and their combinations. C. Number of species studied for each abiotic factor and their combination. Figure made using RAWGraphs (Mauri *et al*. 2017) and the venn R package (Dusa 2022).

**Figure S2.**
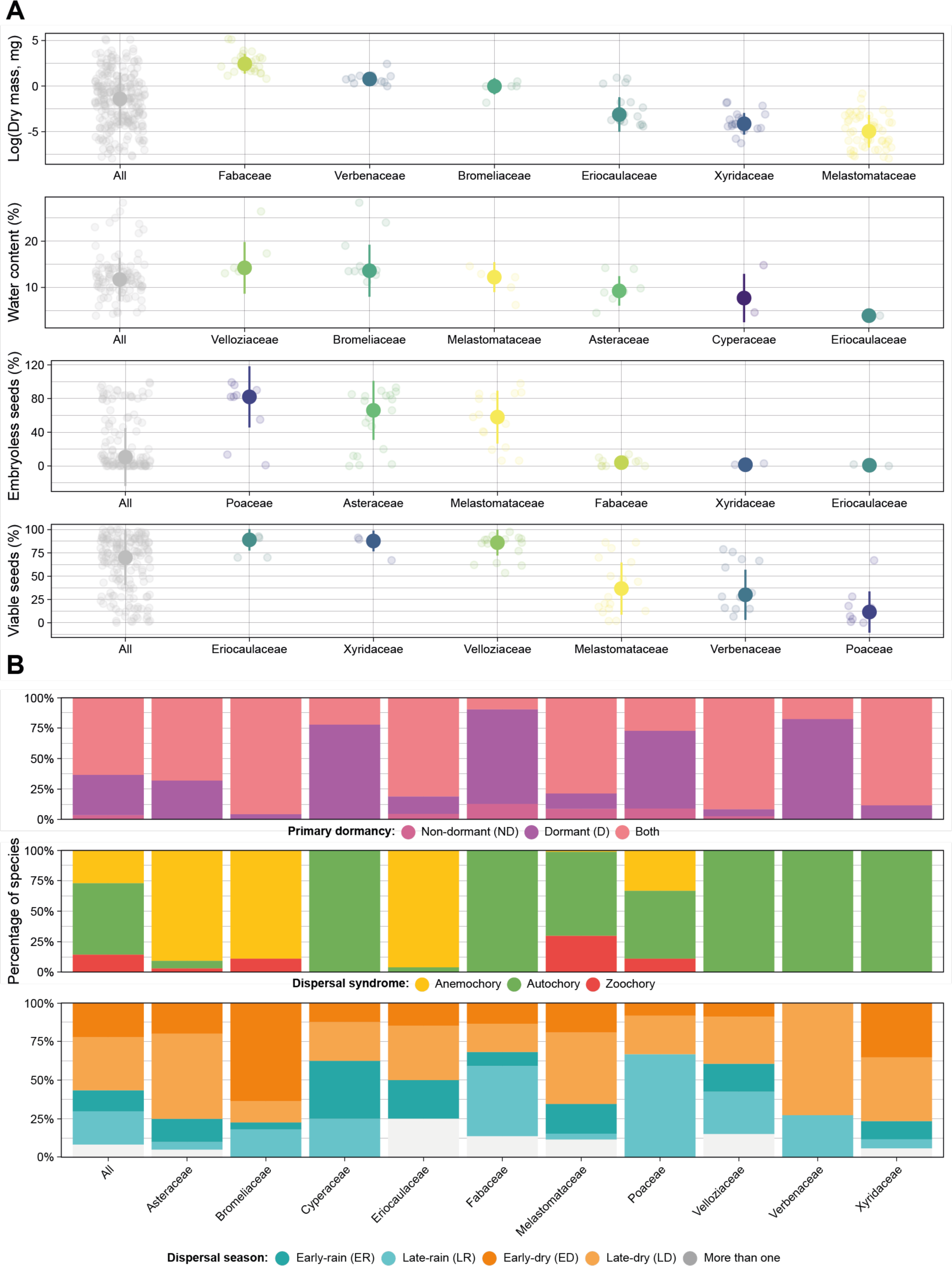
Variation in seven functional seed traits among the most represented families in the database. A. Variation in quantitative traits, showing the variation across the whole dataset and the top three families with the highest and lowest values on each trait. B. Variation in qualitative traits, shown as the proportion of species presenting each trait state across the ten most represented families in the database.

**Table S1.** Moran’s I value for the quantitative seed traits across different taxonomic levels. Bold values indicate values significantly different from zero.

| Seed trait | Genus |  | Family |  | Order |  |
| --- | --- | --- | --- | --- | --- | --- |
|  | I | p.value | I | p.value | I | p.value |
| Seed mass | **0.7035** | **< 0.001** | **0.6545** | **< 0.001** | **-0.3912** | **< 0.001** |
| Water content | **0.4456** | **< 0.001** | **0.2368** | **0.0287** | -0.0623 | 0.4592 |
| Percentage of embryoless seeds | **0.6269** | **< 0.001** | **0.4324** | **< 0.001** | -0.0668 | 0.5038 |
| Percentage of viable seeds | **0.4808** | **< 0.001** | **0.2176** | **< 0.001** | -0.0998 | 0.1337 |

**Table S2.** Summary of phylogenetic mixed models with Bayesian estimation (MCMCglmms) examining the effect of the different abiotic factors, seed mass and dormancy on the final seed germination proportions. Fixed and random effects (±95% CI) and phylogenetic signal λ (±95% CI) are presented. Significant effects are in bold. On average, models were the results of 9000 effect samples.

| Abiotic factor / Predictor | Fixed effects | | | Phylogenetic signal ($\lambda$) | Random effects | | |
| --- | --- | --- | --- | --- | --- | --- | --- |
|  | Posterior mean | 95% C.I. | pMCMC |  | Species | Study | Observation:Studies |
| **Light** |  |  |  |  |  |  |  |
| All | 3.412 | (3.215 ; 3.613) | < **0.001** | 0.884 (0.788 – 0.931) | 7.11 (3.807 – 10.73) | 2.963 (0.456 – 6.542) | 2.114 (1.568 – 2.628) |
| Herb | 2.489 | (2.216 ; 2.758) | < **0.001** | 0.939 (0.843 – 0.973) | 12.99 (4.508 – 24.3) | 10.01 (0.361 – 32.32) | 2.559 (1.925 – 3.223) |
| Shrub | 4.400 | (4.111 ; 4.676) | < **0.001** | 0.859 (0.739 – 0.929) | 6.208 (2.731 – 10.2) | 0.772 (0 – 2.855) | 1.11 (0.53 – 1.71) |
| Restricted | 3.581 | (3.226 ; 3.914) | < **0.001** | 0.893 (0.802 – 0.968) | 9.931 (3.277 – 18.81) | 4.374 (0 – 11.9) | 1.508 (0.79 – 2.193) |
| Widespread | 3.318 | (3.075 ; 3.56) | < **0.001** | 0.869 (0.761 – 0.934) | 6.44 (3.061 – 10.35) | 2.509 (0.185 – 7.074) | 2.327 (1.745 – 2.948) |
| Mesic | 5.282 | (4.906 ; 5.701) | < **0.001** | 0.831 (0.498 – 0.937) | 3.956 (0.457 – 9.199) | 22.59 (0.348 – 71.42) | 1.745 (1.081 – 2.47) |
| Mesic/Xeric | 0.844 | (0.518 ; 1.136) | < **0.001** | 0.946 (0.733 – 0.996) | 16.55 (1.311 – 50.46) | 26.24 (0.033 – 92.11) | 0.944 (0.28 – 1.572) |
| Xeric | 3.448 | (3.159 ; 3.733) | < **0.001** | 0.893 (0.797 – 0.95) | 8.204 (3.61 – 13.74) | 3.652 (0.002 – 9.413) | 1.829 (1.226 – 2.483) |
| Dry mass | -0.437 | (-0.63 ; -0.256) | < **0.001** | 0.846 (0.739 – 0.924) | 5.636 (2.83 – 9.193) | 4.472 (0.614 – 10.45) | 2.099 (1.551 – 2.638) |
| **25/15 °C** |  |  |  |  |  |  |  |
| Overall | -1.384 | (-1.852 ; -0.87) | < **0.001** | 0.917 (0.269 – 0.99) | 5.582 (0.004 – 18.78) | 9.212 (0 – 33.01) | 0.088 (0 – 0.321) |
| Seed mass | -1.259 | (-2.337 ; -0.138) | 0.140 | 0.968 (0.48 – 0.999) | 14.43 (0.049 – 49.4) | 1094 (0 – 1690) | 0.064 (0 – 0.245) |
| **30/20 °C** |  |  |  |  |  |  |  |
| Overall | -0.379 | (-0.828 ; 0.06) | 0.090 | 0.829 (0.491 – 0.983) | 4.018 (0.22 – 11.2) | 8.492 (0 – 29.25) | 0.035 (0 – 0.136) |
| Herb | 0.057 | (-0.746 ; 0.908) | 0.893 | 0.994 (0.007 – 1) | 58.14 (0 – 185.6) | 682.7 (0 – 1462) | 0.093 (0 – 0.359) |
| Shurb | -0.617 | (-1.179 ; -0.038) | **0.036** | 0.931 (0.621 – 0.997) | 13.3 (0.297 – 44.31) | 90.82 (0 – 245.7) | 0.061 (0 – 0.23) |
| Seed mass | 0.322 | (-0.229 ; 0.862) | 0.250 | 0.98 (0.096 – 1) | 16.63 (0 – 60.55) | 776.3 (0 – 1480) | 0.043 (0 – 0.165) |
| **10 °C** |  |  |  |  |  |  |  |
| All | -4.295 | (-4.96 ; -3.613) | < **0.001** | 0.003 (0 – 0.744) | 0.726 (0 – 2.951) | 95.73 (0 – 290.8) | 0.36 (0 – 1.018) |
| Restricted | -2.142 | (-3.194 ; -1.145) | < **0.001** | 0.996 (0.17 – 1) | 183.1 (0 – 640.3) | 140 (0 – 567.9) | 0.35 (0 – 1.314) |
| Widespread | -5.505 | (-6.393 ; -4.691) | < **0.001** | 0.004 (0 – 0.844) | 1.469 (0 – 5.12) | 0 (0 – 0) | 0.187 (0 – 0.645) |
| Dry mass | 1.242 | (0.631 ; 1.843) | < **0.001** | 0.004 (0 – 0.725) | 0.686 (0 – 2.645) | 208.9 (0 – 767.8) | 0.255 (0 – 0.791) |
| **15 °C** |  |  |  |  |  |  |  |
| All | -1.919 | (-2.176 ; -1.663) | < **0.001** | 0.93 (0.876 – 0.969) | 14.12 (6.587 – 22.23) | 1.742 (0.062 – 4.906) | 0.366 (0 – 0.777) |
| Herb | -1.119 | (-1.475 ; -0.786) | < **0.001** | 0.934 (0.852 – 0.98) | 15.2 (3.604 – 31.35) | 13.16 (0.09 – 48.44) | 0.167 (0 – 0.493) |
| Shrub | -2.661 | (-3.045 ; -2.261) | < **0.001** | 0.94 (0.883 – 0.977) | 17.55 (6.621 – 30.81) | 1.407 (0 – 5.84) | 0.506 (0 – 1.047) |
| Restricted | -1.808 | (-2.267 ; -1.363) | < **0.001** | 0.955 (0.849 – 0.986) | 17.2 (3.531 – 38.63) | 2.111 (0 – 8.446) | 0.216 (0 – 0.652) |
| Widespread | -1.975 | (-2.313 ; -1.667) | < **0.001** | 0.943 (0.87 – 0.972) | 15.14 (5.954 – 26.12) | 5.065 (0.089 – 16.77) | 0.507 (0 – 0.993) |
| Mesic | -1.612 | (-1.988 ; -1.262) | < **0.001** | 0.954 (0.868 – 0.983) | 17.18 (4.223 – 34.99) | 16.03 (0.006 – 58.79) | 0.187 (0 – 0.548) |
| Mesic/Xeric | -0.683 | (-1.25 ; -0.159) | **0.015** | 0.973 (0.699 – 0.999) | 20.47 (0.114 – 65.64) | 28.41 (0.017 – 107) | 0.22 (0 – 0.703) |
| Xeric | -2.877 | (-3.346 ; -2.43) | < **0.001** | 0.932 (0.817 – 0.974) | 12.25 (3.305 – 24.24) | 1.601 (0 – 6.554) | 0.583 (0 – 1.223) |
| Dry mass | -0.023 | (-0.309 ; 0.27) | 0.886 | 0.937 (0.872 – 0.968) | 14.15 (6.584 – 23.48) | 2.484 (0.105 – 7.488) | 0.304 (0 – 0.696) |
| **20 °C** |  |  |  |  |  |  |  |
| All | -0.019 | (-0.195 ; 0.165) | 0.838 | 0.843 (0.719 – 0.904) | 4.586 (2.058 – 7.342) | 2.565 (0.15 – 6.241) | 0.119 (0 – 0.348) |
| Herb | 0.666 | (0.395 ; 0.937) | < **0.001** | 0.798 (0.606 – 0.923) | 3.953 (0.952 – 7.842) | 7.608 (0.084 – 24.08) | 0.12 (0 – 0.38) |
| Shrub | -0.572 | (-0.799 ; -0.34) | < **0.001** | 0.902 (0.789 – 0.953) | 7.058 (2.502 – 12.73) | 2.736 (0 – 8.938) | 0.072 (0 – 0.242) |
| Restricted | 0.238 | (-0.088 ; 0.551) | 0.135 | 0.905 (0.699 – 0.962) | 6.047 (1.354 – 13.35) | 3.92 (0 – 10.87) | 0.093 (0 – 0.313) |
| Widespread | -0.148 | (-0.378 ; 0.073) | 0.196 | 0.818 (0.684 – 0.91) | 4.458 (1.82 – 7.669) | 2.979 (0.084 – 8.612) | 0.154 (0 – 0.431) |
| Mesic | 0.095 | (-0.18 ; 0.363) | 0.499 | 0.809 (0.608 – 0.934) | 3.226 (0.803 – 6.923) | 6.351 (0 – 20.27) | 0.035 (0 – 0.129) |
| Mesic/Xeric | 0.463 | (0.038 ; 0.912) | **0.038** | 0.786 (0.486 – 0.956) | 4.477 (0.352 – 11.2) | 6.456 (0 – 21.13) | 0.456 (0 – 1.016) |
| Xeric | -0.386 | (-0.679 ; -0.089) | **0.015** | 0.885 (0.737 – 0.953) | 7.27 (1.926 – 14.07) | 3.559 (0 – 10.35) | 0.205 (0 – 0.561) |
| Dry mass | -0.341 | (-0.534 ; -0.148) | **0.001** | 0.853 (0.741 – 0.916) | 5.059 (2.226 – 8.205) | 3.05 (0.171 – 7.698) | 0.098 (0 – 0.304) |
| **30 °C** |  |  |  |  |  |  |  |
| All | -0.528 | (-0.71 ; -0.342) | < **0.001** | 0.886 (0.816 – 0.937) | 7.243 (3.766 – 11.29) | 1.852 (0.207 – 4.696) | 0.099 (0 – 0.309) |
| Herb | 0.092 | (-0.205 ; 0.357) | 0.518 | 0.685 (0.465 – 0.885) | 2.563 (0.542 – 5.404) | 6.221 (0.138 – 19.95) | 0.167 (0 – 0.476) |
| Shrub | -1.054 | (-1.286 ; -0.819) | < **0.001** | 0.948 (0.893 – 0.971) | 12.4 (5.656 – 20.78) | 1.224 (0 – 4.99) | 0.046 (0 – 0.17) |
| Restricted | -0.264 | (-0.538 ; 0.019) | 0.064 | 0.896 (0.766 – 0.967) | 6.223 (1.507 – 12.92) | 2.957 (0 – 8.832) | 0.03 (0 – 0.117) |
| Widespread | -0.663 | (-0.92 ; -0.418) | 0.196 | 0.904 (0.823 – 0.957) | 9.655 (4.027 – 16.34) | 3.798 (0.112 – 11.08) | 0.223 (0 – 0.563) |
| Mesic | -0.720 | (-0.995 ; -0.435) | < **0.001** | 0.909 (0.802 – 0.971) | 8.003 (2.23 – 16.42) | 9.546 (0 – 34.04) | 0.045 (0 – 0.166) |
| Mesic/Xeric | 0.270 | (-0.206 ; 0.747) | 0.267 | 0.746 (0.393 – 0.962) | 3.905 (0.263 – 10.9) | 9.202 (0.02 – 31.67) | 0.599 (0 – 1.24) |
| Xeric | -0.756 | (-1.032 ; -0.482) | **0.015** | 8.702 (3.05 – 15.57) | 1.852 (0 – 6.214) | 1.852 (0 – 6.214) | 0.085 (0 – 0.289) |
| Dry mass | -0.188 | (-0.388 ; 0.002) | 0.062 | 0.896 (0.815 – 0.941) | 7.612 (3.713 – 12.08) | 2.138 (0.235 – 5.423) | 0.109 (0 – 0.329) |
| **35 °C** |  |  |  |  |  |  |  |
| All | -2.486 | (-2.793 ; -2.196) | < **0.001** | 0.876 (0.717 – 0.928) | 5.625 (2.135 – 9.439) | 4.358 (0.186 – 10.91) | 1.547 (0.903 – 2.191) |
| Herb | -2.153 | (-2.626 ; -1.684) | < **0.001** | 0.845 (0.594 – 0.94) | 4.69 (0.925 – 9.977) | 4.553 (0.067 – 14.7) | 2.243 (1.354 – 3.185) |
| Shrub | -2.725 | (-3.114 ; -2.367) | < **0.001** | 0.858 (0.711 – 0.955) | 7.014 (1.709 – 13.72) | 8.105 (0 – 23.78) | 0.912 (0.142 – 1.585) |
| Restricted | -2.132 | (-2.617 ; -1.673) | < **0.001** | 0.887 (0.686 – 0.966) | 7.348 (1.292 – 15.73) | 2.644 (0 – 8.515) | 0.935 (0 – 1.636) |
| Widespread | -2.660 | (-3.04 ; -2.271) | < **0.001** | 0.854 (0.678 – 0.938) | 5.621 (1.662 – 10.61) | 7.73 (0.086 – 22.35) | 1.891 (1.169 – 2.705) |
| Mesic | -4.019 | (-4.537 ; -3.524) | < **0.001** | 0.784 (0.294 – 0.944) | 3.024 (0 – 7.521) | 31.31 (2.09 – 90.38) | 1.012 (0.173 – 1.904) |
| Mesic/Xeric | -0.929 | (-1.425 ; -0.416) | < **0.001** | 0.889 (0.697 – 0.976) | 8.994 (1.407 – 21.31) | 7.368 (0 – 28.39) | 0.817 (0 – 1.512) |
| Xeric | -2.202 | (-2.661 ; -1.738) | < **0.001** | 0.85 (0.576 – 0.948) | 4.891 (0.716 – 10.62) | 3.076 (0 – 10.06) | 1.597 (0.73 – 2.458) |
| Dry mass | 0.455 | (0.149 ; 0.75) | **0.003** | 0.855 (0.715 – 0.928) | 5.918 (2.302 – 10.32) | 3.504 (0.184 – 9.553) | 1.519 (0.895 – 2.188) |
| **40 °C** |  |  |  |  |  |  |  |
| All | -1.249 | (-1.784 ; -0.708) | < **0.001** | 0.719 (0.398 – 0.87) | 2.332 (0.436 – 5.065) | 0 (0 – 0) | 1.505 (0.538 – 2.446) |
| Herb | -1.188 | (-1.893 ; -0.454) | **0.002** | 0.509 (0.167 – 0.908) | 1.937 (0 – 5.676) | 0 (0 – 0) | 1.224 (0 – 2.392) |
| Shrub | -1.317 | (-2.122 ; -0.524) | **0.001** | 0.798 (0.315 – 0.955) | 3.256 (0.139 – 9.99) | 770.8 (0 – 1331) | 1.845 (0.573 – 3.308) |
| Restricted | -0.915 | (-1.487 ; -0.351) | **0.001** | 0.731 (0.34 – 0.96) | 3.738 (0.138 – 11.52) | 0 (0 – 0) | 0.263 (0 – 0.785) |
| Widespread | -1.629 | (-2.629 ; -0.628) | **0.002** | 0.789 (0.394 – 0.961) | 3.837 (0.107 – 10.45) | 491.2 (0 – 1288) | 3.625 (1.643 – 5.935) |
| Mesic | -3.678 | (-4.699 ; -2.687) | **< 0.001** | 0.993 (0.102 – 1) | 31.33 (0 – 76.51) | 0 (0 – 0) | 0.242 (0 – 0.908) |
| Mesic/Xeric | -1.304 | (-2.238 ; -0.368) | **0.005** | 0.798 (0.413 – 0.992) | 6.474 (0.122 – 20.61) | 509.6 (0 – 1281) | 2.053 (0.399 – 3.835) |
| Xeric | -0.085 | (-0.804 ; 0.657) | 0.809 | 0.744 (0.346 – 0.974) | 3.797 (0.138 – 11.37) | 0 (0 – 0) | 0.938 (0 – 2) |
| Dry mass | -1.149 | (-1.672 ; -0.605) | **< 0.001** | 0.738 (0.436 – 0.902) | 2.708 (0.539 – 6.155) | 579.9 (0 – 1055) | 1.156 (0.277 – 2.051) |
| **Fire-related cues** |  |  |  |  |  |  |  |
| All | -0.641 | (-0.876 ; -0.388) | **< 0.001** | 0.959 (0.887 – 0.982) | 19.54 (6.709 – 36.63) | 1.062 (0 – 3.823) | 0.523 (0 – 0.953) |
| Combined | -2.143 | (-3.46 ; -0.794) | **0.003** | 0.992 (0.832 – 0.999) | 60.61 (0.658 – 200.7) | 0 (0 – 0) | 2.608 (0.458 – 5.382) |
| Single | -0.548 | (-0.784 ; -0.314) | **< 0.001** | 0.952 (0.887 – 0.98) | 18.59 (6.779 – 35.54) | 1.021 (0 – 3.556) | 0.253 (0 – 0.608) |
| Heat | -0.831 | (-1.176 ; -0.485) | **< 0.001** | 0.958 (0.894 – 0.986) | 22.35 (6.755 – 43.45) | 1.455 (0 – 5.276) | 0.496 (0 – 0.992) |
| Smoke | 0.057 | (-0.237 ; 0.342) | 0.697 | 15.44 (0.903 – 0.989) | 15.44 (3.908 – 30.98) | 1.439 (0 – 4.972) | 0.023 (0 – 0.087) |
| Dormant | 0.218 | (-0.248 ; 0.7) | 0.359 | 0.97 (0.487 – 0.999) | 18.91 (0 – 61.13) | 39.82 (0 – 151.1) | 0.059 (0 – 0.223) |
| Non-dormant | -0.814 | (-1.077 ; -0.52) | **< 0.001** | 0.969 (0.901 – 0.992) | 26.69 (6.285 – 56.7) | 2.055 (0 – 6.22) | 0.745 (0 – 1.204) |
| Seed mass | 0.179 | (-0.093 ; 0.454) | 0.198 | 0.958 (0.895 – 0.984) | 21.79 (7.043 – 41.86) | 1.113 (0 – 3.797) | 0.603 (0 – 1.053) |
| **Heat shock temperature** |  |  |  |  |  |  |  |
| 80 °C for 2' | 0.179 | (-0.543 ; 0.903) | 0.616 | 0.964 (0.896 – 0.989) | 18.13 (3.958 – 37.41) | 0.88 (0 – 3.235) | 0.039 (0 – 0.147) |
| 80 °C for 5' | 0.109 | (-0.567 ; 0.798) | 0.755 | 0.953 (0.857 – 0.987) | 15.32 (2.624 – 32.7) | 0.933 (0 – 3.074) | 0.05 (0 – 0.186) |
| 100 °C for 1' | -0.545 | (-1.039 ; -0.089) | **0.026** | 0.963 (0.916 – 0.988) | 19.55 (5.07 – 37.53) | 0.492 (0 – 1.806) | 0.039 (0 – 0.148) |
| 100 °C for 5' | -1.122 | (-1.6 ; -0.663) | **< 0.001** | 0.957 (0.841 – 0.986) | 17.69 (2.814 – 38.09) | 2.62 (0.036 – 8.24) | 0.222 (0 – 0.629) |
| 200 °C for 1' | -2.485 | (-3.096 ; -1.889) | **< 0.001** | 0.964 (0.905 – 0.989) | 21.26 (5.75 – 41.66) | 1.231 (0 – 4.557) | 0.094 (0 – 0.335) |
| **Heat shocks** |  |  |  |  |  |  |  |
| All | -0.500 | (-0.814 ; -0.188) | **0.003** | 0.965 (0.904 – 0.986) | 22.69 (7.102 – 45.39) | 1.13 (0 – 3.766) | 0.17 (0 – 0.494) |
| Herb | -0.373 | (-0.864 ; 0.111) | 0.134 | 0.974 (0.815 – 0.997) | 21.75 (1.228 – 65.16) | 27.61 (0 – 95.44) | 0.064 (0 – 0.245) |
| Shrub | -0.566 | (-0.975 ; -0.157) | **0.006** | 0.98 (0.932 – 0.996) | 42.58 (9.428 – 98.09) | 3.572 (0.041 – 11.95) | 0.312 (0 – 0.788) |
| Restricted | -0.645 | (-1.151 ; -0.177) | **0.007** | 0.925 (0.718 – 0.992) | 12.45 (0.952 – 34.1) | 4.567 (0 – 12.98) | 0.315 (0 – 0.843) |
| Widespread | -0.366 | (-0.789 ; 0.064) | 0.088 | 0.98 (0.929 – 0.995) | 40 (7.305 – 92.64) | 2.755 (0 – 9.573) | 0.126 (0 – 0.416) |
| Mesic/Xeric | -0.072 | (-0.629 ; 0.472) | 0.794 | 0.979 (0.709 – 0.999) | 21.32 (0.231 – 69.83) | 48.92 (0 – 165.4) | 0.045 (0 – 0.173) |
| Xeric | -0.561 | (-0.978 ; -0.148) | **0.007** | 0.974 (0.915 – 0.995) | 37.06 (5.006 – 84.35) | 2.851 (0.001 – 9.76) | 0.383 (0 – 0.895) |
| Dormant | 0.256 | (-0.328 ; 0.898) | 0.406 | 0.983 (0.392 – 1) | 30.67 (0 – 105.5) | 91.08 (0 – 304.2) | 0.08 (0 – 0.305) |
| Non-dormant | -0.722 | (-1.108 ; -0.345) | **< 0.001** | 0.966 (0.914 – 0.993) | 33.35 (6.73 – 74.78) | 1.502 (0 – 4.826) | 0.28 (0 – 0.716) |
| Dry mass | 0.399 | (0.054 ; 0.74) | 0.020 | 0.958 (0.91 – 0.988) | 25.1 (7.582 – 52.74) | 1.321 (0 – 4.325) | 0.18 (0 – 0.52) |
| **Smoke** |  |  |  |  |  |  |  |
| All | 0.057 | (-0.237 ; 0.342) | 0.697 | 0.96 (0.903 – 0.989) | 15.44 (3.908 – 30.98) | 1.439 (0 – 4.972) | 0.023 (0 – 0.087) |
| Herb | 0.446 | (-0.213 ; 1.117) | 0.186 | 0.988 (0.746 – 1) | 35.48 (0.756 – 108.6) | 835.2 (0.135 – 1691) | 0.081 (0 – 0.315) |
| Shrub | -0.025 | (-0.368 ; 0.308) | 0.885 | 0.975 (0.928 – 0.995) | 24.68 (4.67 – 55.35) | 2.393 (0 – 8.081) | 0.023 (0 – 0.087) |
| Restricted | 0.181 | (-0.294 ; 0.662) | 0.468 | 0.924 (0.699 – 0.993) | 10.47 (0.717 – 30.87) | 12.97 (0 – 40.24) | 0.073 (0 – 0.276) |
| Widespread | -0.021 | (-0.398 ; 0.399) | 0.922 | 0.983 (0.928 – 0.998) | 34.03 (3.48 – 86.99) | 2.347 (0 – 8.289) | 0.027 (0 – 0.107) |
| Mesic/Xeric | 0.243 | (-0.556 ; 1.058) | 0.548 | 0.997 (0.56 – 1) | 97.85 (0.046 – 300.5) | 0 (0 – 0) | 0.1 (0 – 0.382) |
| Xeric | 0.018 | (-0.328 ; 0.339) | 0.913 | 0.973 (0.911 – 0.993) | 20.47 (3.877 – 46.81) | 2.13 (0 – 6.92) | 0.024 (0 – 0.092) |
| Dormant | 0.499 | (-0.342 ; 1.37) | 0.254 | 0.995 (0.195 – 1) | 114.7 (0 – 331.1) | 1085 (0 – 1737) | 0.139 (0 – 0.543) |
| Non-dormant | -0.021 | (-0.331 ; 0.299) | 0.895 | 0.972 (0.904 – 0.993) | 19.07 (4.138 – 42.55) | 1.601 (0 – 5.143) | 0.027 (0 – 0.105) |
| Dry mass | 0.000 | (-0.317 ; 0.332) | 0.996 | 0.973 (0.915 – 0.992) | 19.81 (4.747 – 42.28) | 1.802 (0 – 6.374) | 0.027 (0 – 0.104) |

**Table S3.** Summary of phylogenetic mixed models with Bayesian estimation (MCMCglmms) examining the effect of the different abiotic factors, seed 1 mass and dormancy on median germination time (t_50_). Fixed and random effects (±95% CI) and phylogenetic signal λ (±95% CI) are presented. Significant effects are in bold. On average, models were the results of 9000 effect samples.

| Abiotic factor / Predictor | Fixed effects | | | Phylogenetic signal ($\lambda$) | Random effects | | |
| --- | --- | --- | --- | --- | --- | --- | --- |
|  | Posterior mean | 95% C.I. | pMCMC |  | Species | Study | Observation:Studies |
| 25/15 °C |  |  |  |  |  |  |  |
| Overall | 6.620 | (5.385 ; 7.917) | **0.001** | 0.998 (0.993 – 1) | 426.1 (85.92 – 976.1) | 157.4 (0 – 662.7) | 6.288 (3.909 – 9.031) |
| Seed mass | 5.732 | (3.162 ; 8.24) | **< 0.001** | 0.997 (0.975 – 1) | 265.4 (13.37 – 803) | 2758 (0 – 5097) | 4.239 (2.078 – 6.755) |
| 30/20 °C |  |  |  |  |  |  |  |
| Overall | 1.234 | (0.524 ; 1.956) | **0.002** | 0.993 (0.974 – 0.999) | 132.9 (22.82 – 317.9) | 104.5 (0 – 415.7) | 1.772 (0.76 – 2.842) |
| Herb | 2.241 | (1.196 ; 3.298) | **< 0.001** | 0.998 (0.852 – 1) | 284.8 (1.744 – 844.1) | 1672 (0 – 2427) | 0.906 (0 – 2.209) |
| Shurb | 0.734 | (-0.216 ; 1.679) | 0.123 | 0.996 (0.98 – 1) | 235 (19.35 – 644.4) | 341.9 (0 – 1163) | 2.117 (0.781 – 3.554) |
| Seed mass | 1.003 | (0.149 ; 1.862) | **0.027** | 0.997 (0.98 – 1) | 236.5 (23.78 – 632.5) | 3957 (0 – 2924) | 1.657 (0.537 – 2.916) |
| 15 °C |  |  |  |  |  |  |  |
| All | 5.419 | (4.691 ; 6.15) | **< 0.001** | 0.99 (0.976 – 0.997) | 98.21 (32.69 – 185.4) | 19.19 (0 – 61.37) | 6.87 (5.289 – 8.482) |
| Herb | 5.125 | (4.385 ; 5.875) | **< 0.001** | 0.982 (0.949 – 0.994) | 140 (22 – 337.4) | 253.9 (0.009 – 625.7) | 4.116 (2.859 – 5.552) |
| Shrub | 5.903 | (4.399 ; 7.479) | **< 0.001** | 0.992 (0.973 – 0.998) | 116.2 (24.07 – 262.7) | 18.35 (0 – 75.27) | 11.99 (8.108 – 16.47) |
| Restricted | 7.089 | (5.617 ; 8.432) | **< 0.001** | 0.995 (0.92 – 1) | 114.9 (0 – 357) | 51 (0 – 186.6) | 7.286 (4.296 – 10.57) |
| Widespread | 4.752 | (3.89 ; 5.598) | **< 0.001** | 0.992 (0.973 – 0.998) | 115.1 (32.65 – 232.5) | 28.61 (0 – 114.8) | 6.52 (4.81 – 8.425) |
| Mesic | 5.018 | (3.351 ; 6.654) | **< 0.001** | 0.995 (0.969 – 1) | 174.8 (11.05 – 491.2) | 98.63 (0 – 388.8) | 10.7 (6.57 – 15.07) |
| Mesic/Xeric | 6.509 | (5.503 ; 7.565) | **< 0.001** | 0.993 (0.941 – 0.999) | 106.8 (5.34 – 320.2) | 108.5 (0 – 329) | 4.348 (2.577 – 6.308) |
| Xeric | 4.674 | (3.482 ; 5.873) | **< 0.001** | 0.99 (0.94 – 0.999) | 92.89 (2.359 – 262.6) | 50.7 (0 – 167.2) | 6.444 (4.055 – 9.138) |
| Dry mass | -1.109 | (-1.842 ; -0.311) | 0.886 | 0.992 (0.974 – 0.997) | 102.4 (27.17 – 206) | 30.66 (0 – 99.35) | 6.684 (5.052 – 8.398) |
| 20 °C |  |  |  |  |  |  |  |
| All | 1.037 | (0.512 ; 1.549) | **< 0.001** | 0.997 (0.994 – 0.999) | 343.7 (171.4 – 542.8) | 11.86 (0.001 – 41.13) | 6.5 (5.318 – 7.678) |
| Herb | 0.835 | (0.279 ; 1.395) | **0.003** | 0.982 (0.949 – 0.994) | 49.8 (13.3 – 101.3) | 51.14 (0.206 – 165.7) | 2.709 (1.792 – 3.618) |
| Shrub | 1.200 | (0.362 ; 2.029) | **0.005** | 0.982 (0.998 – 0.999) | 495.2 (192.7 – 855.8) | 32.46 (0 – 125.6) | 9.707 (7.555 – 11.98) |
| Restricted | 0.796 | (-0.067 ; 1.644) | 0.066 | 0.994 (0.985 – 0.998) | 165 (56.15 – 314.2) | 18.19 (0 – 65.24) | 6.619 (4.831 – 8.509) |
| Widespread | 1.187 | (0.529 ; 1.86) | **0.002** | 0.998 (0.995 – 0.999) | 495.3 (170.9 – 892.5) | 39.31 (0 – 157.9) | 6.446 (5.038 – 7.993) |
| Mesic | 1.185 | (0.579 ; 1.82) | **< 0.001** | 0.999 (0.998 – 1) | 1622 (491.7 – 3199) | 181.6 (0.087 – 732.7) | 2.188 (1.203 – 3.156) |
| Mesic/Xeric | 1.494 | (0.196 ; 2.805) | **0.029** | 0.984 (0.92 – 0.998) | 52.5 (4.734 – 137) | 61.51 (0 – 209.5) | 9.704 (6.574 – 12.96) |
| Herb | -0.410 | (-0.869 ; 0.052) | 0.082 | 0.982 (0.949 – 0.994) | 43.9 (10.39 – 93.8) | 82.06 (0.165 – 287.6) | 1.497 (0.834 – 2.277) |
| Shrub | 1.690 | (0.515 ; 2.841) | **0.005** | 0.995 (0.986 – 0.998) | 179.9 (54.14 – 349.5) | 37.17 (0 – 137.5) | 16.38 (12.76 – 20.3) |
| Restricted | -0.955 | (-2.023 ; 0.132) | 0.083 | 0.993 (0.974 – 0.998) | 106.3 (24.7 – 219.6) | 27.19 (0 – 92.75) | 8.628 (6.18 – 11.32) |
| Widespread | 1.432 | (0.635 ; 2.177) | **< 0.001** | 0.994 (0.984 – 0.998) | 154.8 (48.1 – 291.4) | 44.67 (0 – 181.8) | 8.959 (7.125 – 10.89) |
| Mesic | 2.930 | (1.457 ; 4.459) | **< 0.001** | 0.996 (0.975 – 1) | 194.4 (16.49 – 519.9) | 96.69 (0 – 394.6) | 15.14 (10.62 – 20.12) |
| Mesic/Xeric | -1.394 | (-2.409 ; -0.455) | **0.008** | 0.962 (0.732 – 0.998) | 20.24 (0.168 – 60.27) | 23.59 (0 – 95.38) | 4.378 (2.775 – 6.149) |
| Xeric | 0.315 | (-0.489 ; 1.169) | **0.471** | 0.995 (0.987 – 0.998) | 180.3 (60.2 – 343.4) | 35.32 (0 – 119.1) | 6.96 (5.31 – 8.946) |
| Dry mass | -0.624 | (-1.535 ; 0.254) | 0.062 | 0.995 (0.988 – 0.998) | 181.4 (74.25 – 324.6) | 28.59 (0 – 96.32) | 10.05 (8.306 – 11.97) |
| **35 °C** |  |  |  |  |  |  |  |
| All | 2.151 | (1.417 ; 2.943) | **< 0.001** | 0.994 (0.985 – 0.998) | 148.8 (58.67 – 264) | 28.29 (0 – 90.39) | 9.471 (7.506 – 11.32) |
| Herb | 2.239 | (1.472 ; 2.951) | **< 0.001** | 0.982 (0.949 – 0.994) | 114.9 (24.03 – 246.9) | 54.54 (0.001 – 195.5) | 5.052 (3.67 – 6.525) |
| Shrub | 2.027 | (0.408 ; 3.629) | **< 0.001** | 0.992 (0.966 – 0.997) | 94.29 (17.9 – 215.4) | 73.75 (0 – 230.6) | 16.98 (12.32 – 22.44) |
| Restricted | 0.405 | (-0.893 ; 1.686) | 0.530 | 0.997 (0.988 – 0.999) | 258.3 (50.35 – 589.7) | 54.97 (0 – 213.3) | 8.923 (5.986 – 12.15) |
| Widespread | 3.085 | (2.121 ; 4.017) | **< 0.001** | 0.991 (0.976 – 0.998) | 111.7 (26.43 – 230.7) | 67.71 (0 – 211) | 9.349 (7.114 – 11.57) |
| Mesic | 2.665 | (1.06 ; 4.247) | **< 0.001** | 0.998 (0.985 – 1) | 360.7 (25.81 – 1005) | 98.31 (0 – 385.8) | 7.729 (4.36 – 11.41) |
| Mesic/Xeric | 1.924 | (0.231 ; 3.553) | **0.022** | 0.985 (0.859 – 0.999) | 47.28 (0.001 – 138.1) | 42.77 (0 – 175.6) | 15.76 (11.24 – 21.18) |
| Xeric | 2.173 | (1.262 ; 3.119) | **< 0.001** | 0.988 (0.955 – 0.997) | 74.39 (12.88 – 178.8) | 93.63 (0 – 296.2) | 6.013 (4.089 – 7.908) |
| Dry mass | 0.613 | (-0.193 ; 1.48) | 0.156 | 0.993 (0.983 – 0.998) | 139.4 (47.69 – 259.6) | 41.18 (0 – 125.8) | 10.09 (7.994 – 12.24) |
| **Fire-related cues** |  |  |  |  |  |  |  |
| All | -0.283 | (-0.898 ; 0.296) | 0.349 | 0.994 (0.983 – 0.998) | 159.6 (41.53 – 333.9) | 28.57 (0.296 – 112) | 5.565 (4.357 – 6.777) |
| Combined | -1.885 | (-3.496 ; -0.171) | **0.035** | 0.998 (0.981 – 1) | 720.4 (0.109 – 2327) | 0 (0 – 0) | 1.686 (0 – 3.876) |
| Single | -0.260 | (-0.851 ; 0.402) | 0.413 | 0.993 (0.983 – 0.998) | 158.2 (41.7 – 327.9) | 33.63 (0.288 – 127.9) | 5.82 (4.543 – 7.083) |
| Heat | 0.012 | (-0.925 ; 0.922) | 0.971 | 0.992 (0.973 – 0.999) | 114.6 (19.27 – 250.5) | 45.23 (0.317 – 160.5) | 7.243 (5.256 – 9.338) |
| Smoke | -0.974 | (-1.67 ; -0.31) | **0.004** | 0.997 (0.988 – 0.999) | 274.3 (50.7 – 653.4) | 76.75 (0.141 – 240) | 2.187 (1.189 – 3.27) |
| Dormant | -1.494 | (-3.551 ; 0.586) | 0.157 | 0.996 (0.921 – 1) | 207.3 (207.3 – 0) | 1348 (0 – 3135) | 9.817 (5.092 – 15.53) |
| Non-dormant | -0.088 | (-0.728 ; 0.513) | 0.777 | 0.995 (0.985 – 1) | 232.8 (40.24 – 574) | 77.03 (0.319 – 259.1) | 5.053 (3.916 – 6.274) |
| Seed mass | -0.608 | (-1.239 ; 0.039) | 0.063 | 0.994 (0.981 – 0.999) | 160.2 (35.3 – 353.6) | 34.6 (0.207 – 131.6) | 5.881 (4.627 – 7.205) |
| **Heat shock temperature** |  |  |  |  |  |  |  |
| 80 °C for 2' | -1.463 | (-2.994 ; 0.075) | 0.061 | 0.995 (0.986 – 0.999) | 196.5 (53.11 – 415.4) | 25.71 (0 – 100.5) | 2.91 (1.529 – 4.515) |
| 80 °C for 5' | -0.466 | (-1.765 ; 0.91) | 0.493 | 0.996 (0.988 – 0.999) | 225.5 (58.69 – 483.5) | 36 (0.002 – 141.9) | 3.112 (1.724 – 4.683) |
| 100 °C for 5' | 0.944 | (-0.519 ; 2.229) | 0.173 | 0.993 (0.968 – 0.999) | 123.1 (11.64 – 287.8) | 10.93 (7.717 – 14.52) | 1.046 (0.671 – 1.486) |
| **Heat shocks** |  |  |  |  |  |  |  |
| All | -0.066 | (-1.029 ; 0.836) | 0.893 | 0.992 (0.972 – 0.998) | 110.9 (19.99 – 243) | 44 (0.485 – 162.5) | 7.255 (5.32 – 9.377) |
| Herb | -1.184 | (-2.448 ; 0.111) | 0.070 | 0.993 (0.965 – 0.999) | 130.4 (11.62 – 350.5) | 199.8 (0.342 – 603.7) | 5.474 (3.118 – 8.125) |
| Shrub | 0.716 | (-0.555 ; 2.074) | 0.285 | 0.996 (0.949 – 1) | 0.716 (-0.555 – 2.074) | 190 (0.001 – 609.1) | 66.62 (0 – 221.2) |
| Restricted | -0.900 | (-1.862 ; 0.086) | 0.067 | 0.995 (0.979 – 0.999) | 183.8 (23.73 – 463.2) | 171 (0.296 – 616.9) | 3.893 (2.22 – 5.462) |
| Widespread | 1.017 | (-0.756 ; 2.855) | 0.272 | 0.994 (0.153 – 1) | 87.79 (0 – 346.7) | 41.93 (0.001 – 143) | 12.54 (7.75 – 18.28) |
| Mesic/Xeric | -2.855 | (-4.804 ; -1.027) | **0.005** | 0.997 (0.934 – 1) | 187.6 (0.01 – 607.7) | 1163 (0 – 2761) | 6.932 (2.923 – 11.87) |
| Xeric | 1.043 | (-0.139 ; 2.109) | 0.073 | 0.004 (0 – 0.963) | 6.092 (0 – 25.93) | 27.71 (0.374 – 96.94) | 6.755 (4.596 – 9.351) |
| Dormant | -1.983 | (-3.955 ; 0.006) | 0.052 | 0.996 (0.95 – 1) | 218.2 (0 – 689.2) | 0 (0 – 0) | 7.042 (2.952 – 11.98) |
| Non-dormant | -0.723 | (-1.108 ; -0.345) | **< 0.001** | 0.995 (0.97 – 0.999) | 33.35 (6.73 – 74.78) | 1.502 (0 – 4.826) | 0.28 (0 – 0.716) |
| Dry mass | -0.964 | (-1.987 ; -0.033) | 0.053 | 0.993 (0.96 – 0.998) | 97.35 (10.7 – 233.3) | 51.02 (0.14 – 189.8) | 7.646 (5.45 – 9.844) |
| **Smoke** |  |  |  |  |  |  |  |
| All | -0.974 | (-1.67 ; -0.31) | **0.004** | 0.997 (0.988 – 0.999) | 274.3 (50.7 – 653.4) | 76.75 (0.141 – 240) | 2.187 (1.189 – 3.27) |
| Herb | -1.560 | (-3.548 ; 0.411) | 0.120 | 0.999 (0.976 – 1) | 600.5 (6.639 – 1777) | 1176 (0 – 1618) | 7.226 (3.348 – 12.26) |
| Shrub | -0.785 | (-1.294 ; -0.298) | **< 0.001** | 0.998 (0.99 – 1) | 414.3 (42.7 – 1113) | 58.16 (0.183 – 207) | 0.19 (0 – 0.575) |
| Restricted | -1.447 | (-2.409 ; -0.417) | **0.007** | 0.997 (0.988 – 1) | 359.8 (44.06 – 950.3) | 699.9 (0 – 1312) | 3.595 (1.971 – 5.405) |
| Widespread | -0.071 | (-0.655 ; 0.528) | 0.819 | 0.999 (0.461 – 1) | 907.6 (0.001 – 1348) | 160.5 (0.272 – 243.6) | 0.051 (0 – 0.196) |
| Non-dormant | -1.090 | (-1.665 ; -0.541) | **< 0.001** | 0.997 (0.988 – 0.999) | 295.1 (50.39 – 711.9) | 65.08 (0.299 – 225.1) | 1.026 (0.248 – 1.858) |
| Dry mass | -0.131 | (-0.848 ; 0.622) | 0.724 | 0.997 (0.984 – 0.999) | 265.1 (37.71 – 699.9) | 86.51 (0.11 – 231.4) | 2.349 (1.276 – 3.48) |

